# *Pseudomonas aeruginosa* MipA-MipB envelope proteins act as new sensors of polymyxin

**DOI:** 10.1101/2023.08.14.553335

**Authors:** Manon Janet-Maitre, Viviana Job, Maxime Bour, Sabine Brugière, Mylène Robert-Genthon, David Cobessi, Yohann Couté, Katy Jeannot, Ina Attrée

## Abstract

Due to the rising incidence of antibiotic resistant infections, the last-line antibiotics polymyxins have resurged in the clinics in parallel with new bacterial strategies of escape. The Gram-negative opportunistic pathogen *Pseudomonas aeruginosa* develops resistance to colistin/polymyxin by distinct molecular mechanisms, mostly through modification of the lipid A component of the LPS by proteins encoded within the *arnBCDATEF-ugd* (*arn*) operon. In this work, we characterized a polymyxin-induced operon, named *mipBA*, notably present in *P. aeruginosa* strains devoid of the *arn* operon. We showed that *mipBA* is activated by ParR/ParS two-component regulatory system in response to polymyxin. MipA and MipB localize to bacterial outer membrane and form a complex *in vitro*. Structural modeling revealed that the lipoprotein MipB adopts a β-lactamase fold with two additional C-terminal domains,while MipA folds as an outer-membrane β-barrel, harboring an internal negatively charged channel, able to host a polymyxin molecule. Nano differential scanning fluorimetry (DSF) showed that polymyxin stabilized MipA protein *in vitro*. Mass spectrometry-based quantitative proteomics on whole bacterial membranes demonstrated that the Δ*mipBA* mutant synthesized less MexXY-OprA proteins in response to polymyxin compared to the wild-type strain, as a consequence of impaired transcriptional activation of the *mex* operon. We propose MipA/MipB to act as membrane (co)sensors working in concert to activate ParS histidine kinase and help the bacterium to cope with polymyxin-mediated envelope stress through synthesis of the efflux pomp, MexXY-OprA.

## Introduction

*Pseudomonas aeruginosa* is a Gram-negative opportunistic pathogen, which thrives in a wide range of environments and displays high intrinsic resistance to antibiotics, the latter being one of the main threats to the modern healthcare system. In 2017, the World Health Organization (WHO) classified *P. aeruginosa* as a critical priority pathogen for which the development of novel antibiotics is urgently needed. Polymyxins, a class of antibiotics which includes polymyxin B (PMB) and polymyxin E (PME, also known as colistin), are currently used as a last resort to treat multiresistant *P. aeruginosa* infections [1,2]. Polymyxins are amphipathic cationic antimicrobial peptides (cAMPs), which interact with the negatively charged lipid A component of the lipopolysaccharide (LPS), resulting in its destabilization and loss of outer membrane integrity. Although the exact mechanism of bacterial killing is still elusive, polymyxin insertion in the outer membrane alters membrane curvature and stability [3]. In the proposed model, self-promoted uptake of polymyxins lead to a contact between the inner and outer membranes, allowing phospholipid exchange, in turn creating osmotic imbalance and eventually leading to bacterial cell death [4–6]]. While polymyxins have a strong bactericidal effect on *P. aeruginosa*, the latter can adapt to polymyxin stress by inducing a set of eight genes, named *arnBCADTEF-ugd* (*arn*). As a final product, Arn enzymes synthesize the 4-amino-4-deoxy-L-arabinose (L-Ara4N) moiety that is transferred to a nascent lipid A in the inner membrane by ArnT [7]. This LPS modification reduces the overall negative charge of the outer membrane, decreasing the affinity of polymyxins toward the bacterial surface.

Expression of the *arn* operon is tightly regulated in response to external stimuli by at least four two component regulatory systems (TCS) composed of a membrane sensor histidine kinase (HK) and a response regulator (RR) [8]. PhoP/PhoQ and PmrA/PmrB are able to activate the *arn* locus in response to low magnesium concentration, whereas ParR/ParS and CprR/CprS respond to the presence of different cAMPs including polymyxins through an unknown mechanism [9–11]]. In addition to the *arn* operon, ParR/ParS system down-regulates the expression of the OprD major porin gene, which contributes to carbapenem entry into *P. aeruginosa* and up-regulates the expression of *pmrAB*, *mexXY* operon coding for an efflux pump, and that of the *PA1797* gene, encoding an uncharacterized protein annotated as a putative β-lactamase [9,12]. Amino-acid substitutions in either ParR or ParS were associated with a significant decrease of polymyxin susceptibility in clinical strains of *P. aeruginosa* due to overexpression of the *arn* operon, as well as to carbapenems and MexXY-OprM/A efflux substrates (cefepim, fluoroquinolones, and aminoglycosides) [13–15]].

Recent studies investigating the genetic diversity of *P. aeruginosa* isolates revealed five distinct phylogenetic groups/clades [16,17]. According to the classification from Freschi *et al.*, 2019 [16], the phylogenetic group 3, harboring the fully sequenced strain PA7 [18], was the most distant to the two predominant groups, represented by reference strains PA14 (group 1) and PAO1 (group 2). Comparison of gene content of different groups revealed that all genes encoding the type three secretion system (T3SS) were absent from both groups 3 and 5. These groups encoded a cytolytic two-partner secretion system, ExlB/ExlA [19]. Surprisingly, whereas the gene *arnA* was present in all sequenced strains from other groups, only 38.5% of strains belonging to group 3 harbored the *arnA* gene, raising the question about the origins and evolution of this group of strains, as well as their mechanism of adaptation to polymyxins [16].

In this work, we showed that despite the absence of the *arn* operon, a recent clinical isolate of group 3, called IHMA879472 (IHMA87, [20,21]), is capable of adaptation to polymyxins. In all investigated strains from group 3, the polymyxin-responsive gene *IHMA87_03332/PA1797* (renamed here *mipB*), is encoded in a ParR/ParS-regulated two-gene operon together with *mipA*. We evidenced that MipA and MipB are associated with the outer membrane and interact *in vitro*. Comparative proteomic analysis of envelope proteins following PMB challenge, showed that the deletion of *mipBA* led to a significant decrease in efflux pump proteins MexXY and OprA, known to be regulated at transcriptional level by ParS/ParR. In addition, we showed that MipA can interact with PMB by nano differential scanning fluorimetry (nano-DSF). This data confirms structural predictions by Alphafold showing MipA as a β-barrel outer membrane protein with an internal negatively charged channel able to accommodate the polymyxin B (PMB) molecule. As the *mipBA* deletion abolished *mexXY-oprA* induction in response to PMB, we propose that MipA acts as a co-sensor of ParS-ParR regulatory system. MipA could entrap polymyxins and transmit the signal through its partner MipB, to the inner-membrane HK sensor protein ParS, thus representing a new concept of detection of this class of antibiotics.

## Results

### L-Ara4N is not essential for polymyxin adaptation

Unlike other species such as *Escherichia coli* and *Acinetobacter baumannii*, the addition of L-Ara4N to the lipid A was sufficient to confer polymyxin resistance to both selected *in vitro* mutants and clinical strains of *P. aeruginosa* [22]. The strain IHMA87 lacks the entire *arn* operon encompassing the deletion of approx 8.8 kb between genes *algA* and *fruA*, corresponding to PA3551 and PA3560 in PAO1 (Fig. 1A). To evaluate the frequency of this event, we compared genomes available on the NCBI database belonging to groups 3 (*n=*23), 4 (*n*=64) and 5 (*n*=38) with a set of strains of groups 1 (strain PA14, *n=*8) and 2 (strain PAO1, *n=*7). Interestingly, only a subset of strains of group 3 carried the deletion of the *arn* locus reminiscent to the IHMA87 genome, whereas 31.6% of strains (subgroup 3A, *n*=12, strain PA7) possessed the complete *arn* operon (Fig. 1B).

**Figure 1.**
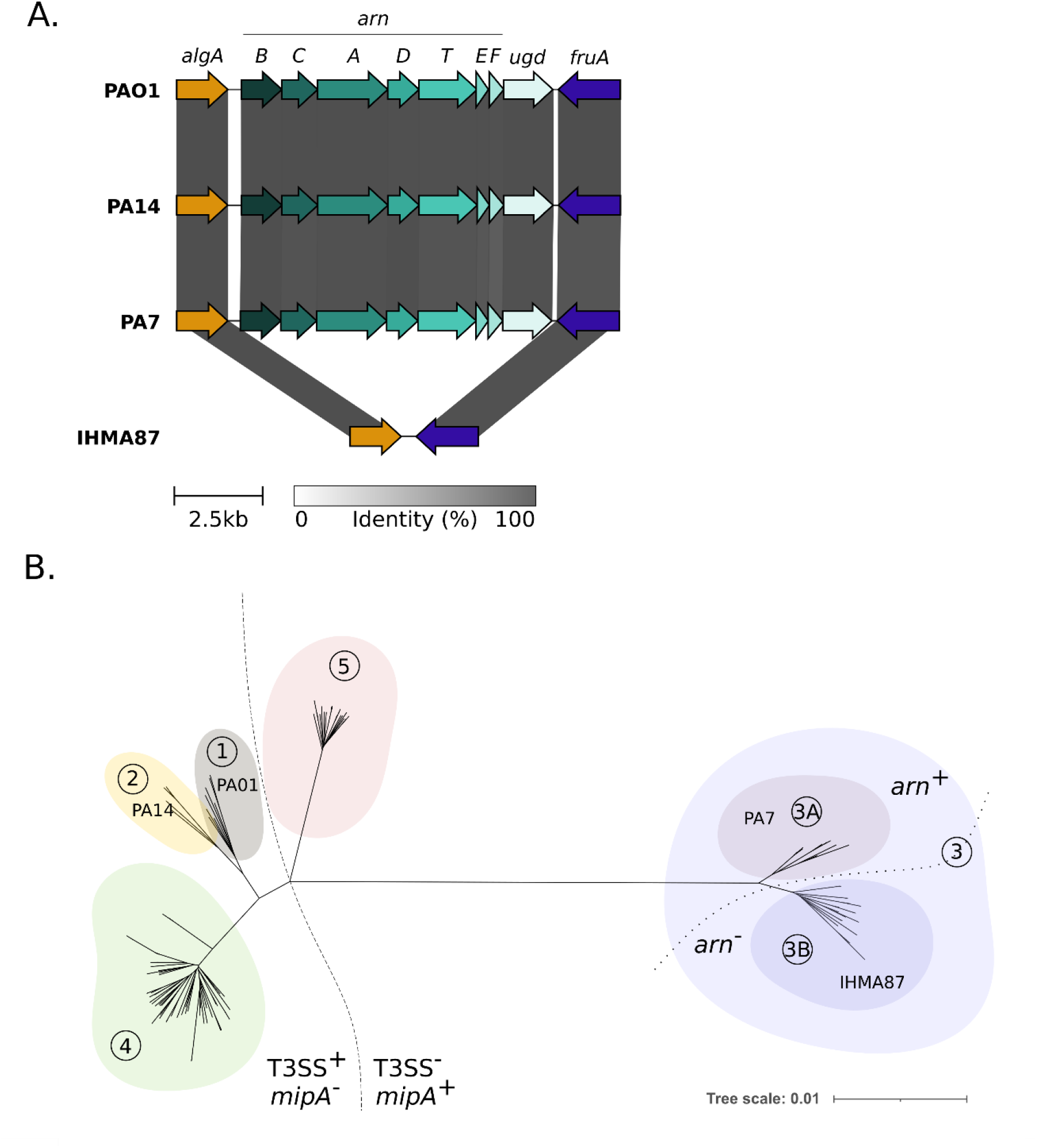
Identification of *P. aeruginosa* strains lacking *arn* operon. A. Conservation of *arn* locus across *P. aeruginosa* strains, visualized by Clinker. The *arn-ugd* region expending from *algA* to *fruA* encompasses 8.8 kb and is not present in IHMA87. **B.** Neighbor Joining phylogenetic tree highlighting separation of phylogenetic group 3 into two distinct subgroups 3A and 3B, which differ notably by the presence of the *arn* locus.

The minimal inhibitory concentration (MIC) of the PME, towards eight selected strains of group 3B, including IHMA87, was identical to those of reference strains PAO1, PA14 and PA7 (MIC=0.5 mg/L) in agreement with a previous study indicating that *arn* operon is not involved in intrinsic polymyxin resistance [23]. Acquired resistance to polymyxins in clinical strains of *P. aeruginosa* are associated with mutations in one or several genes encoding TCS PmrA/PmrB, ParR/ParS, PhoP/PhoQ and CprR/CprS, leading to constitutive *arn* operon expression and L-Ara4N addition to LPS [24,25]. We therefore evaluated the impact of this large deletion on the selection of PME resistant mutants *in vitro*. In contrast to strains PAO1, PA7 and PA14, no resistant mutant was obtained from IHMA87 grown on Mueller Hinton agar plates supplemented with 8 to 64 mg/L of PME, showing that the *arn* operon is required to acquire stable PME resistance. The same was observed with several strains lacking *arn,* ZW26, JT87, CPHL1145, LMG5031. Interestingly, we observed that according to a genetic background, the rates differed from 1.20 10^-7^ (±5.92 10^-8^) to 6.67 10^-9^ (±6.66 10^-9^) (Fig. 2A).

**Figure 2.**
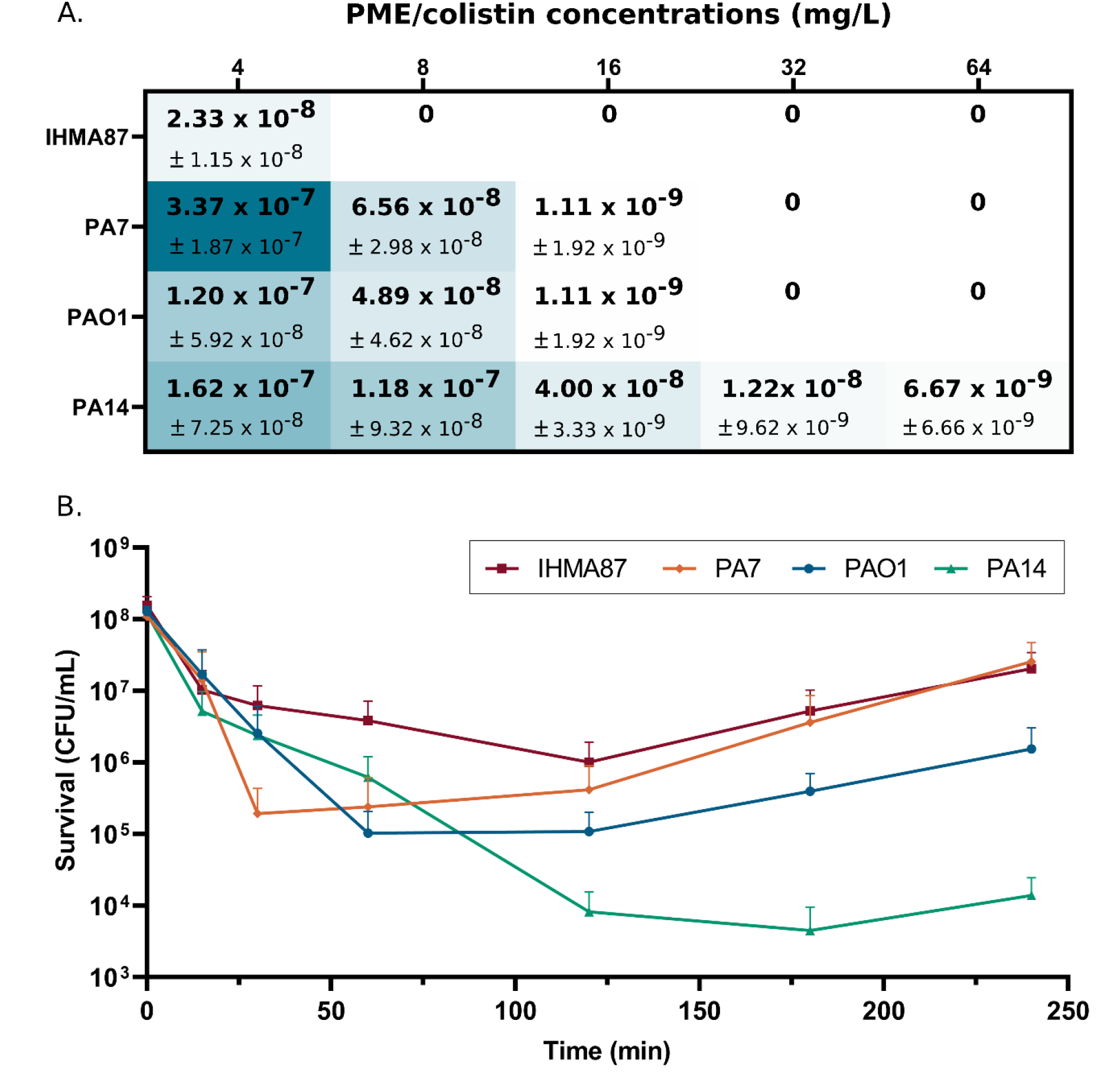
Arn operon is necessary for acquisition of stable resistance but not for adaptive resistance to polymyxins. **A** Mutation rates in IHMA87 and reference strains PA7, PAO1 and PA14 in presence of 4 to 64 mg/L PME. **B.** Bactericidal activity of polymyxins on IHMA87, PA7, PAO1 and PA14 strains over time in presence of 8xMIC (4mg/L) of PME. n=3.

*P. aeruginosa* is also able to tolerate polymyxins in growth medium through the induction of three TCS PmrA/PmrB, ParR/ParS, and CprR/CprS [9,12,26–28]. To test the tolerance of the arn negative strain, IHMA87, the strains were exposed to step-by-step increase concentration of PME. Interestingly, as the others, the IHMA87 strain was able to grow with up to 64 mg/L of PME, but with a lower number of viable bacteria (Fig. S1A; 2.2 ×10^4^ +/-7.7×10^4^ CFU/mL *versus* 6.7 ×10^6^+/-1.1×10^6^ CFU/mL for strain PAO1, 8.3 ×10^4^+/-3.1×10^4^ CFU/mL for strain PA14, and 2.4 ×10^7^+/-3.1×10^7^ CFU/mL for strain PA7) whereas growth in absence of polymyxin was comparable (Figure S1B). Overall, this data suggests that the capacity of adaptation to PME persists in the IHMA87. To confirm that IHMA87 was able to respond to PME, we determined the bactericidal effect of supra concentrations (8x-MIC) of PME. As indicated in Fig. 2B, a higher bactericidal effect was observed for the strains PAO1 and PA14 in comparison to strains IHMA87 and PA7 after 30 minutes post exposure. The regrowth started after 60 and 120 minutes was more pronounced for PA7 and IHMA87 leading to a 1-3 log difference in bacterial CFU after 240min of treatment. The absence of L-Ara4N synthesis does not seem to alter the adaptation of the strain IHMA87 to PMB. Overall, a strain devoid of *arn* operon is able to adapt to polymyxin stress, suggesting alternative *arn*-independent mechanisms in play.

### *mipBA* is activated by polymyxins in a ParR/ParS-dependent manner

In order to investigate the possible role of other genes in PMB adaptation, we focused on the gene of unknown function *IHMA87_03332/PA1797* (renamed *mipB*), which is induced by PMB in a ParR/ParS-dependent manner [9,12]. The genetic environment of *mipB* across *P. aeruginosa* strains is variable (Fig. 3A). Although the syntheny of the *parRS* operon and mipB is conserved across the clades, strains of group 3 harbor a second gene immediately downstream of *mipB* annotated as *mipA* for MltA-interacting protein A (Fig. 3A). In PA14 and PAO1 the predicted open reading frames of residual *mipA* is 117 nucleotides-long compared to the 780 bp *mipA* gene in IHMA87 and PA7. The residual predicted protein MipA* shares more than 60% identity over 38-amino acid-long sequences, suggesting a genetic remodeling of the region and partial loss of *mipA* sequences. tRNA coding sequences, frequently found as hotspots for genetic remodeling [29,30], are present just downstream of *mipA* (*Pseudomonas* genome database; [31]).

**Figure 3.**
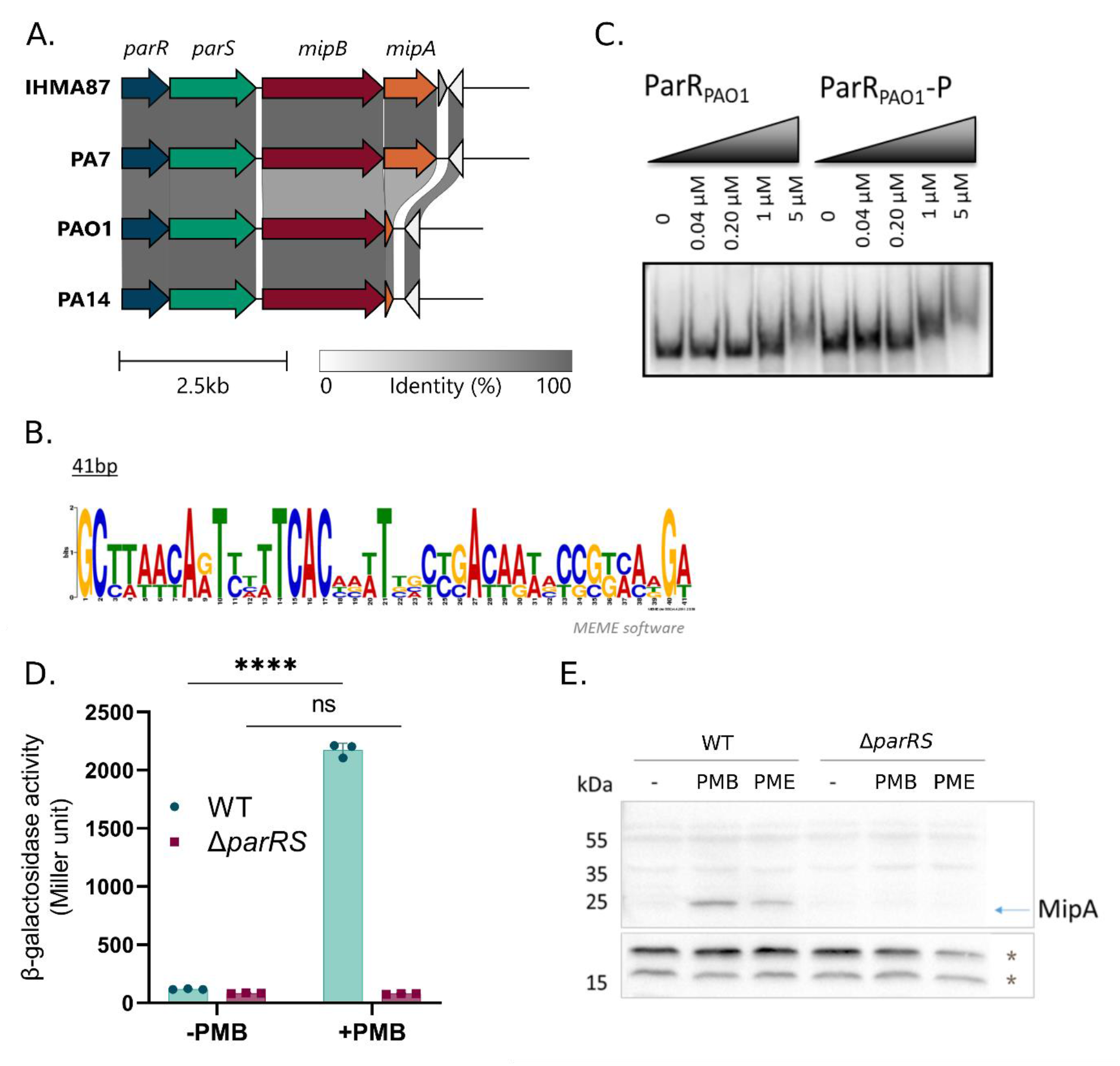
MipA is synthetized in response to PMB in a ParR/ParS-dependent manner. **A** Conservation of *mipBA* locus across *P. aeruginosa* strains, visualized by Clinker. **B**. Consensus of ParR binding site obtained from known target promoters**. C.** EMSA showing ParR binding onto the promoter of *PA1797* with or without phosphorylation (P). **D.** *P_mipBA_* activity in response to PMB measured by β-galactosidase assay in wild type (IHMA87) and IHMA87Δ*parRS* mutant. **E.** MipA detection in response to sub-lethal concentrations of PMB and PME in a ParRS-dependent manner. * Indicates nonspecific antibody binding used as loading control.

As the ParR regulon has been largely investigated [9,12,32], the ParR binding consensus sequence using the bioinformatics tool MEME could be defined (Fig. 3B). A perfect ParR binding site, conserved across the reference strains, was identified upstream the ATG of *mipB*. We examined whether the regulation of *mipB* is direct using purified ParR protein and electrophoresis mobility shift assay (EMSA) on a predicted promoter region of *PA1797* (PAO1). In agreement with the recent genome-wide binding pattern of ParR [32], we showed a direct binding of ParR to the promoter of *PA1797*, which is improved when ParR is phosphorylated (Fig. 3C). We then used RT-qPCR to re-examine the expression and predicted operonic structure of *mipBA* in IHMA87, in presence of PMB. The polycistronic RNA *mipB-mipA* was produced in high amounts in response to PMB (30-fold increase), confirming that the two genes form an operon (Fig. S2A). To confirm the role of ParR in *mipBA* gene expression, we designed a transcriptional reporter and measured *P_mipBA_* promoter activity in response to a sub-lethal dose of PMB (0.25 µg/mL). As shown in Figure 3D, *P_mipBA_* activity was increased 18-fold in response to a sub-lethal PMB treatment in a ParR/ParS-dependent manner. A dose-response effect of PMB on *P_mipBA_* activity was observed increasing up to 0.3 µg/mL PMB before reaching saturation (Fig. S2B).

We then examined the levels of MipA synthesis in response to PMB using specific polyclonal antibodies. MipA was detected in response to sub-lethal PMB treatment and was undetectable in a *parRS* deletion mutant (Fig. 3E). Using MipA-directed antibodies and a version of MipB with 3xFLAG tag at the C-terminus (MipB_3xFLAG_), we followed the kinetic of MipB and MipA production upon the addition of a sub-lethal concentration of PMB. MipB_3xFLAG_ was detected from 15 min after the addition of PMB (Fig. S2C) and the quantities of MipB_3xFLAG_ increased up to 60-90 min, respectively. MipB was not detected in late stationary phase. Low amounts of MipA were detected 30 min after PMB addition, reached a maximum at 90 min and was still detectable in late stationary phase. These results show that both MipB and MipA are rapidly produced in response to PMB treatment and reached their maximum abundance between 60 and 120 min, with MipB being less stable. To determine whether PMB-dependent induction of MipA was conserved in other T3SS-negative strains, we used nine isolates from group 3 and 5 ([33], Fig. S2D). Overall, MipA induction in response to sub-lethal concentration of PMB appears to be a conserved mechanism except for the strain DVL1758, in which the protein could not be detected. Interestingly, in one strain, Zw26, MipA synthesis was constitutively produced, which probably results from a constitutive activity of the ParS/ParR system.

Overall, the ParR-dependent production of MipA and MipB is triggered by a sub-lethal concentration of PMB and PME and is conserved across representative strains from the groups 3 and 5 of *P. aeruginosa*.

### MipA and MipB are associated with the bacterial membranes

MipA and MipB are predicted to localize to the bacterial envelope. MipB carries a type II signal peptide (residues M1-G21) and an “ASGC” lipobox sequence characteristic of bacterial lipoproteins (Fig. 4A and Fig. S3). The amino-acids at positions +2 and +3 after the Cys, suggest that MipB is targeted to the outer membrane [34,35]. MipB harbors a β-lactamase fold but lacks the “SXXK” motif necessary for activity (Fig. 4B, [36]). MipB also shows degenerated motif 2 (YXN) and 3 (KTG), present in the *E. coli* K12 β-lactamase AmpC (Fig. 4B, [37,38]). In addition, MipB possesses 220 additional residues at its C-terminus that are absent in *E. coli* AmpC, suggesting this domain may provide specific function to *P. aeruginosa* MipB.

**Figure 4.**
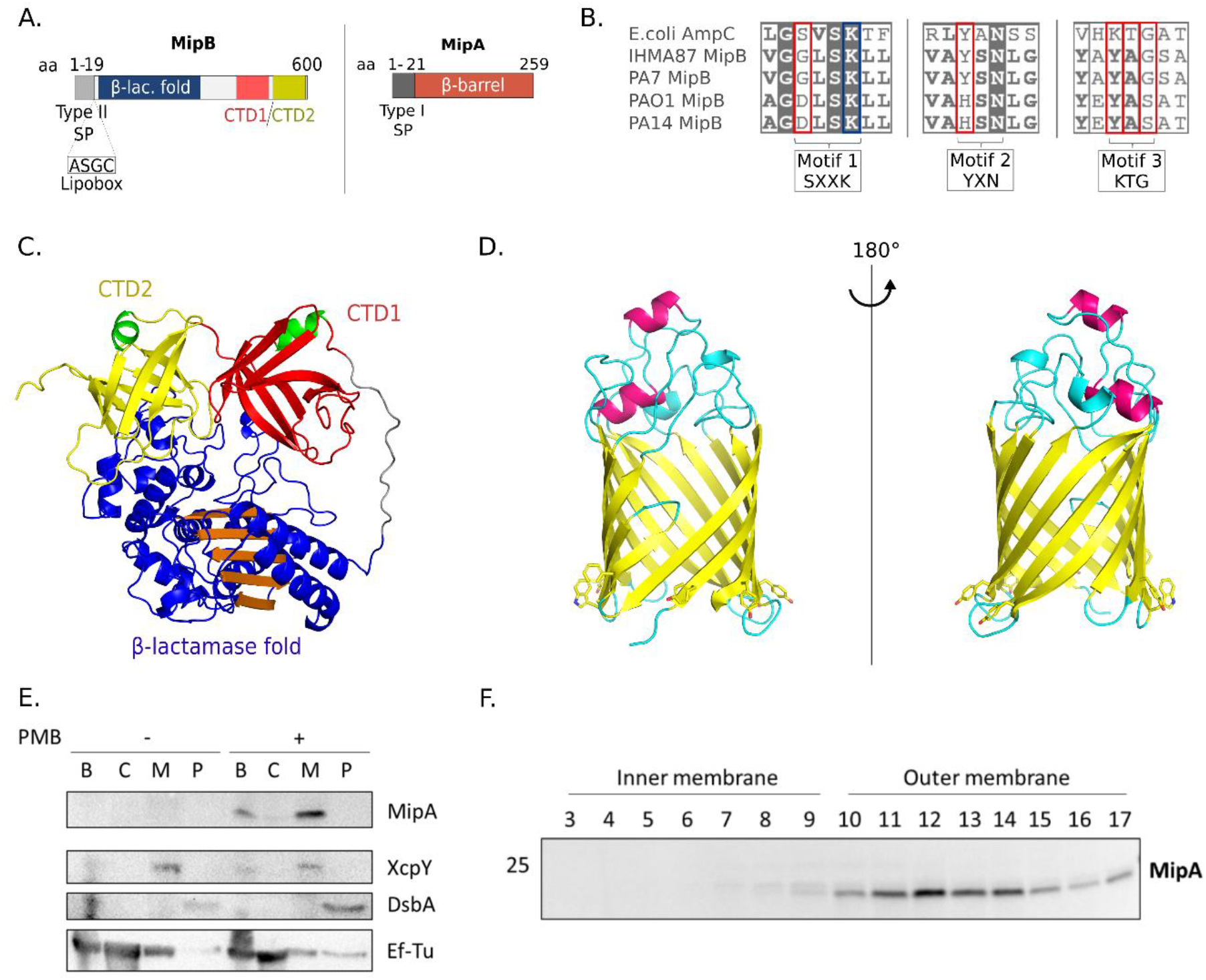
MipB and MipA are envelope proteins. **A** Schematic representation of MipA and MipB. **B.** Alignment of the catalytic sites of AmpC (*E. coli*) with MipB from different *P. aeruginosa* strains. **C.** MipB model calculated using AlphaFold excluding the 27 first residues containing the “lipobox”, which anchors the protein to the membrane, and last 8 residues. The β-strands are represented as arrows and α-helices as ribbons. The N-terminal domain containing the β-lactamase fold is in blue and orange. The two eight-stranded antiparallel β-barrels in C-terminal are colored in red and yellow, and their short N-terminal α-helices in green. The long loop connecting the C-terminal to the N-terminal domain is in gray. **D.** MipA model generated by AlphaFold, without the 21 first residues containing the signal peptide (Met1-Ala21). The β-strands are represented as yellow arrows and α-helices as pink ribbons. The side chains of aromatic residues that delineate the inner membrane are drawn in sticks. Figures were generated with Pymol. **E-F.** Experimental validation of the model. Bacterial fractionation showing membrane association of MipA (**E.**). EF-Tu, XcpY and DsbA are controls for cytosol, membrane, and periplasmic fractions, respectively. Lines B: whole bacteria, C: cytosol, M: total membranes, P: periplasm. **F.** Inner and outer membrane separation by sucrose gradient showing the presence of MipA in the outer membrane fractions.

Using AlphaFold [39], we generated a model for MipB and retrieved the MipA model (Fig. 4C-D). As suggested by sequence alignment (Fig. S3), MipB harbors a “serine” β-lactamase-like fold (residues M28-S387) with a central five-stranded antiparallel β-sheet surrounded by α-helices similar to the structure of β-lactamases and penicillin binding proteins (PBPs; [40,41]). Two additional C-terminal domains, CTD1 (V405-I501) and CTD2 (L506-R593), folded as an eight-stranded antiparallel β-barrel, are connected to the main domain by a linker formed by a long loop (G388-A404). Both are closed by a short N-terminal α-helix (Fig. 4C). Interestingly CTD1 and CTD2 are conserved in MipB proteins in strains of three clades independently of the presence of MipA (Fig. S3). These two domains have folds similar to the C-terminal domain of penicillin binding protein Rv0907 from *Mycobacterium tubercolosis* H37RV (PDB 4WHI; [42]) of unknown function. The analysis of the electrostatic potential of these two domains shows a large positively charged surface exposed to the external part of the protein.

MipA, belonging to the MipA/OmpV family, harbors a N-terminal signal peptide (M1-A21) predicted to be cleaved according to SignalP [43], and folds as a β-barrel with 12 transmembrane β strands. The presence of five tryptophan residues (four of them highly conserved in MipA proteins, Fig. S4) suggests that the β-barrel membrane insertion occurs *via* the same mechanism observed for OmpA, without the need of Bam complex (Fig. 4D; [44]). The 12 β-strands are connected by long extracellular loops that close the lumen of the barrel, and periplasmic turns given the localization of the N- and C-termini on the same side [44]. The lumen of the barrel is opened on the periplasmic space and on the side through a lateral gate delineating a channel that connects the periplasm to extracellular space. The β-barrel has a cross-section of ∼ 25 Å×22 Å and is ∼40 Å in height. Using PDBeFold [45] and FoldSeek [46], MipA structure superimposes on several structures from PDB such as OmpG (PDB entry: 2X9K; [47]), and NanC (PDB entries: 2WJR, 2WJQ; [48]) with root-mean-square deviation (rmsd) values of 2.55 and 2.58 Å, two porins from *E. coli* involved in the N-acetylneuraminic and maltodextrin transport, respectively. MipA also superimposes onto the barrel domain of the intimin protein from *E. coli* (PDB entry: 5G26; [49]) and the lipid A deacylase LpxR of *Salmonella typhimurium* (PDB entry: 3FID; [50]) with rmsd values of 2.33 and 3.01 Å, respectively. Using FoldSeek, several predicted structures by AlphaFold as β-barrels with 12 transmembrane β-strands and the lateral gate superimpose onto MipA; these proteins belonging to different species of Gram-negative bacteria.

In order to experimentally corroborate the predictions, we used a bacterial fractionation method to separate the cytoplasm, periplasm and membranes, and investigate protein localization in bacteria following PMB challenge. The immunoblotting performed on different fractions confirmed the membrane localization of both MipA and MipB (Fig. 4E and Fig. S5). We further separated the inner and outer membranes on a sucrose gradient [51,52], and confirmed that MipA is associated with the outer membrane (Fig. 4F). Finally, both proteins MipA and MipB were found associated with bacterial membranes (Fig. 4E-F and S5), showing that both proteins are tightly associated to the bacterial membrane, in line with the structural predictions.

To investigate on a potential interaction between the two proteins, a complex between MipA and MipB was built using AlphaFold-Multimer [53]. Its analysis reveals two main interaction zones that encompass the eight-stranded antiparallel β-barrels of the MipB CTDs that interact with the open-periplasmic side of MipA (Fig. 5A), mainly through charged residues. To confirm the relevance of this structural model, we co-produced MipB and MipA in *E. coli* and performed an affinity chromatography experiment. Proteins were expressed in *E. coli* from a bicistronic vector yielding MipA-His_6_ and MipB-Strep. Due to their membrane localization, we tested the effective solubility of both proteins in the presence of different detergents. The highest solubility was obtained in the presence of N-lauroylsarcosine. MipB-Strep and its potential partner were purified by affinity chromatography on a Strep-column. The obtained fractions were analyzed by SDS-PAGE followed by Coomassie staining or Western blotting. The eluted fraction contained both MipA and MipB, suggesting the complex formation (Fig. 5B).

**Figure 5.**
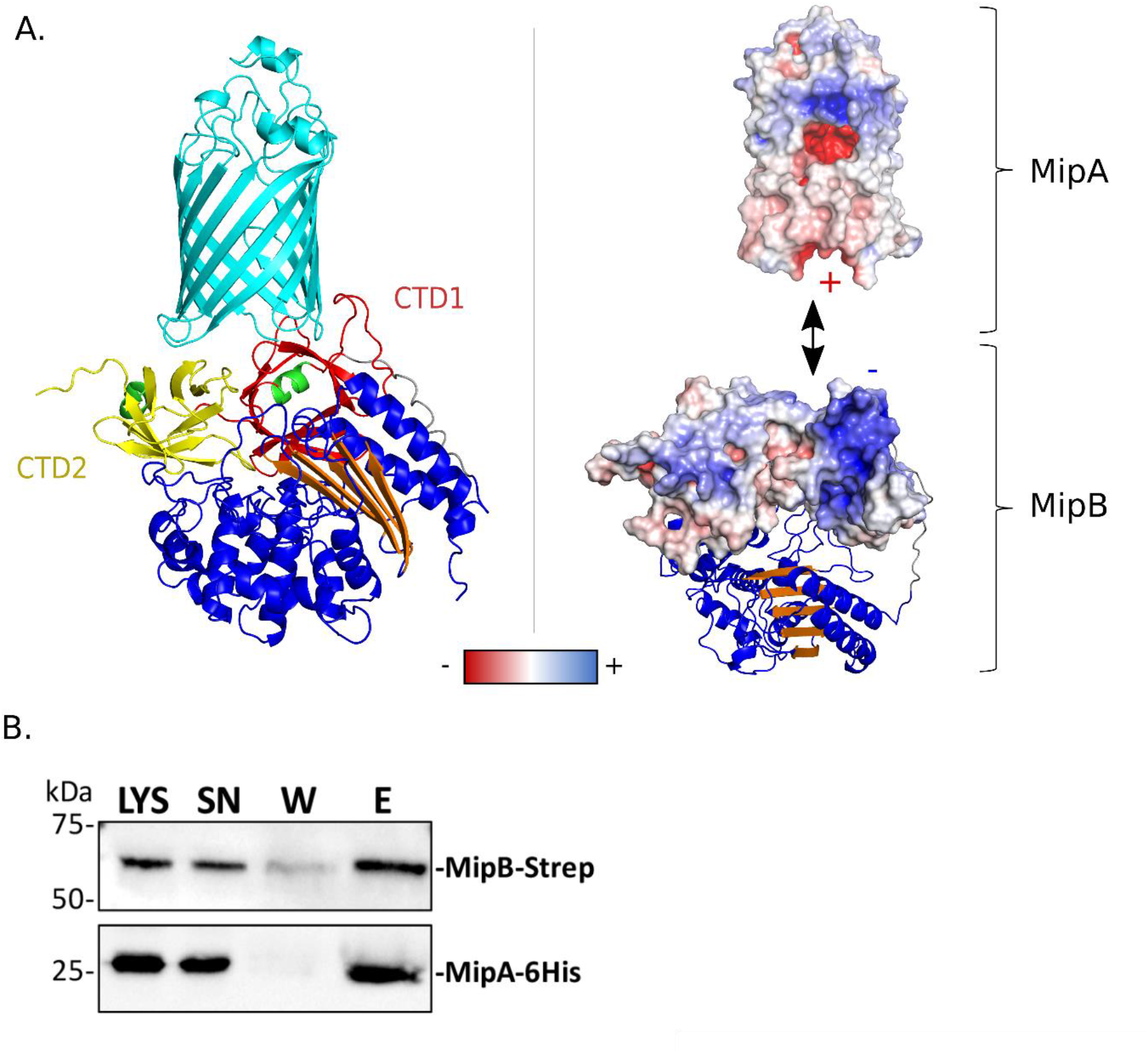
MipB and MipA complex. **A** Model of the MipA-MipB complex generated using AlphaFold-Multimer (Evans *et al*., 2022). Left: MipA is colored in cyan. The β-lactamase fold of MipB is in blue and orange. The two 8 anti-parallel β-stranded barrel domains of MipB are colored in yellow and red with the first α-helices in green. Right: Electrostatic surface representation showing the charges at the interface between the two proteins. The figures were generated with Pymol. **B.** MipB and MipA co-purify *in vitro*. MipB-Strep and MipA-His_6_ were co-produced in *E. coli*. Soluble extracts were added to a Strep column and proteins were eluted by the addition of a desthobiotin containing buffer. Different fractions were analyzed by immunoblotting using anti-Strep and anti-MipA primary antibodies.

Taken together, both MipB and MipA are membrane-associated proteins and form a complex *in vitro*.

### MipA binds PME and PMB

The electrostatic surface analyzes of MipA shows that the protein is hydrophobic on the outside of the β-barrel, the hydrophobic membrane part being delineated by a girdle of aromatic residues at the periplasmic interface, coherent with a membrane-embedded protein. Interestingly, the channel of MipA is mainly composed of negatively charged amino acids (Fig. 6A, left) that together with its size are compatible for binding of positively charged molecules such as PMB. We therefore tested this hypothesis by docking PMB and PME into MipA using Autodock Vina [54] and found solutions (Fig. 6A, right), suggesting that MipA may bind PMB and PME.

**Figure 6.**
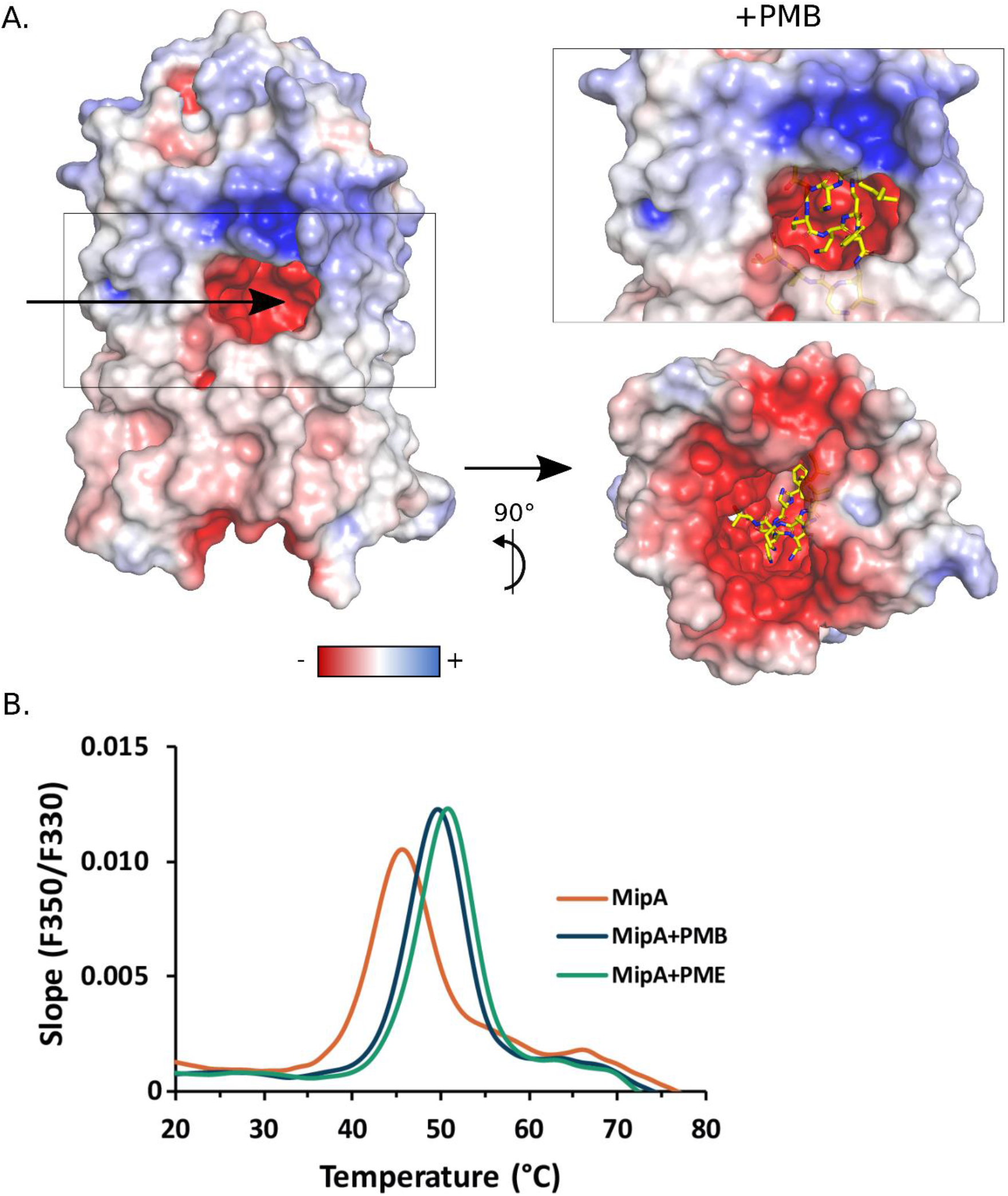
MipA binds PMB and PME. **A.** PMB docking in MipA. Left: MipA electrostatic surface. The arrow shows the negative channel entrance. Right top: Membrane perpendicular view from the PMB bound to MipA. The best position of PMB calculated with Autodock Vina [54] is shown in sticks: the carbon, nitrogen and oxygen atoms are colored in yellow, blue and red respectively. Right bottom: View from the MipA periplasmic face of the PMB bound to MipA. **B**. Thermal stability of MipA increases in presence of PMB and PME. Pure MipA alone (in orange) or incubated for 2h at RT with a molar ratio of (1:4) PMB (dark blue) or PME (in green) were heated from 20 to 95°C. Protein unfolding was followed by Tryptophan fluorescence at 330 and 350nm using nano-DSF technology. The slope of the ratio (F350/F330) is plotted at the different temperatures, the maximum corresponds to the melting temperature of the protein.

To experimentally validate the model, we performed nano-DSF to determine protein stability in presence of cAMPs employing intrinsic tryptophan and tyrosine fluorescence. Purified MipA (Fig. S6A) was mixed with PMB and PME at different molar ratio (MipA:ligand) from 1:1 to 1:4. Samples were incubated at room temperature for 2h and then analyzed by nano-DSF (Nanotemper) to determine the apparent melting temperature (Tm) as a measure of protein folding and stability. In the purification buffer containing 0.1% LAPAO, MipA displayed a Tm of 45.6°C. Upon addition of PMB and PME, the Tm values increased to 49.6°C and 50.8°C, respectively (Fig. 6B), but not in presence of the buffer alone (Fig. S6B), inferring conformational changes in MipA. A similar shift in Tm of 6°C was observed previously upon PMB binding to LSD1 demethylase [55].

These results show a stabilization of MipA in presence of both AMPs and suggest a direct binding of the two molecules to MipA.

### *mipBA* deletion alters bacterial response to polymyxins

Taking into account the envelop localization of MipA/MipB and polymyxin binding to MipA, we hypothesized that MipA/MipB work in concert to defend the bacterial cell against polymyxin-induced envelope damage. We adopted a mass spectrometry-based quantitative proteomic analysis of bacterial membranes to investigate bacterial response to polymyxin B through the comparison of protein abundances in a Δ*mipBA* mutant compared to the parental strain (Table S1). In the wild-type IHMA87 strain, treatment with a sub-lethal concentration of polymyxin B led to the drastic increase of both MipA and MipB in bacterial membranes (Fig. 7A), in agreement with our results showing induction of the operon by PMB, on the transcriptional level and increased levels of both proteins observed by Western blot. Interestingly, MipA but not MipB was detected in the membranes of the wild-type strain in basal condition, showing a residual amount of MipA present at all times in the membranes (undetectable by Western blot; Fig. 3E). In addition to Mip proteins, the quantities of the tripartite efflux pump MexXY-OprA (IHMA87_03098-IHMA87_03100) were increased (with Log_2_(FC) of 3-5.45), in line with *mexXY-oprA* overexpression upon PMB treatment in other strains [9,12].

**Figure 7.**
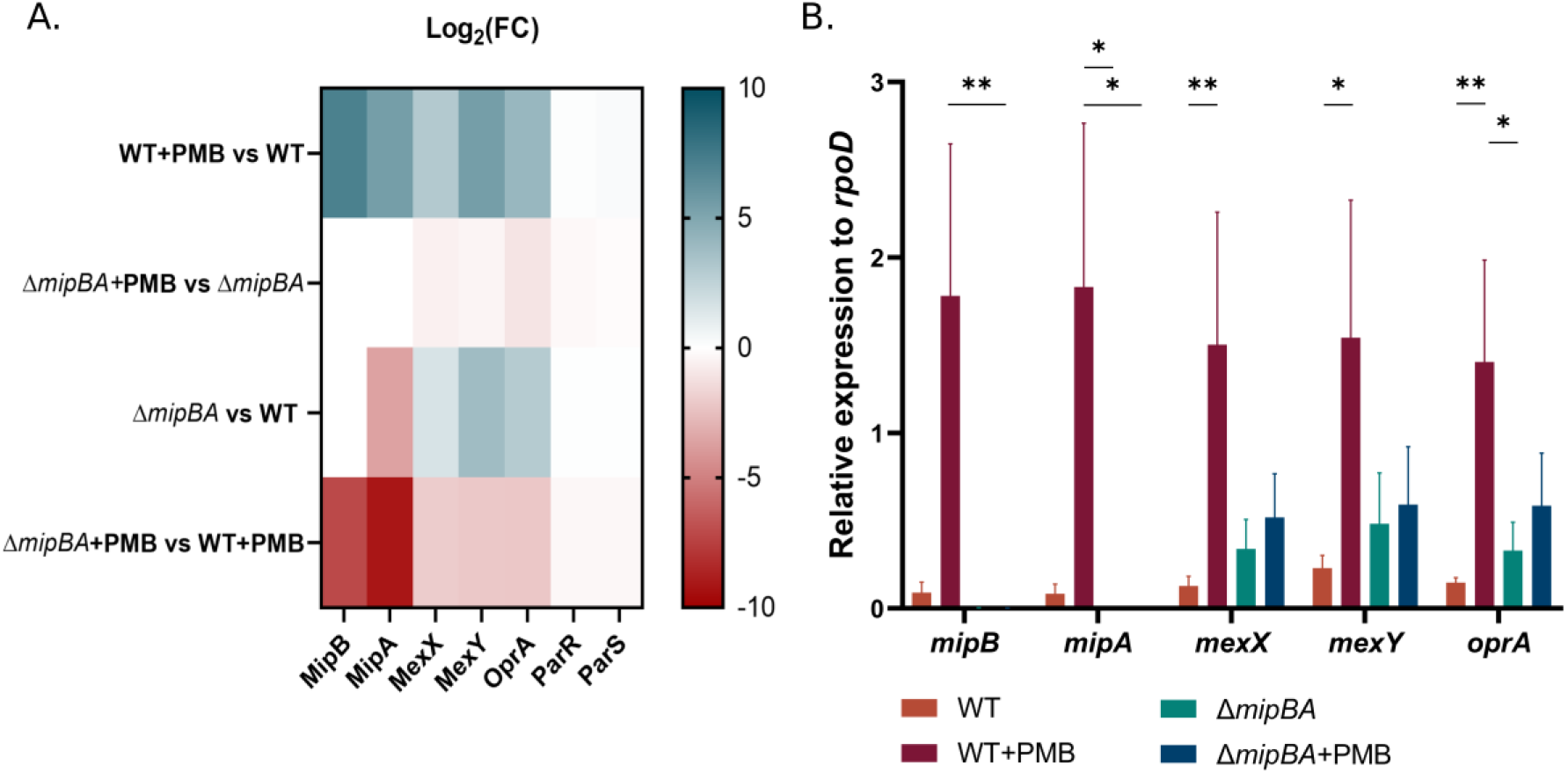
MipB and MipA are required for *P. aeruginosa* response to PMB. **A** Proteomic analysis of IHMA87 membranes with or without sub-lethal PMB treatment. The quantity of proteins overexpressed in the wild-type strain upon PMB treatment (MipA, MipB, MexX, MexY and OprA) was compared in different conditions. All off-white boxes were statistically significant. **B.** mRNA quantities (relative expression normalized to *rpoD*) of *mipB, mipA, mexX, mexY* and *oprA* in IHMA87 (WT) or Δ*mipBA* with or without addition of sub-lethal concentrations of PMB. A Kruskal-Wallis test was applied followed by a Dunn’s test for each of the genes tested.

The comparison of membrane proteomes upon PMB treatment showed significant decrease of the three proteins of the MexXY-OprA system in the membranes of Δ*mipBA* (Log_2_(FC) ranging between −2.02 to −2.21; Fig 7A). This is of interest, as *mexXY* transcriptional activation in presence of PMB depends on ParR/ParS signaling [9,12], suggesting that MipA and/or MipB could assist ParR/ParS in the sensing of polymyxins to activate the downstream signaling pathway leading to the induction of ParR regulon responsible for the adaptive resistance to polymyxins. To note, the quantities of ParR and ParS were identical in the two conditions, and similar in the two strains (wild type versus Δ*mipBA*), in agreement with previous studies showing that PMB does not activate the *parRS* operon at the transcriptional level [9].

As the response of *mexXY-oprA* operon to PMB occurs at the transcription level, we validated our data using RT-qPCR on bacterial cultures challenged with PMB (Fig. 7B). Indeed, the three genes of the operon *mexXY-oprA* in IHMA87 were activated by PMB ranging from 6.46- to 11.64-fold increase. In the *mipBA* mutant, this upregulation was significantly decreased. Interestingly, although non ignificant, higher levels of *mex* transcripts could be measured in the *mipBA* mutants in non-activated conditions (without PMB), in agreement with the comparative proteomics data. It is tempting to speculate that in the absence of the activator (here PMB), the low levels of MipA in the membranes, detected in proteomics (Table S1) of the wild-type strain maintain ParS/ParR system in inactive form, keeping the ParR regulon shut-down.

Overall, the absence of MipB/MipA led to a dysregulation of the PMB adaptive response, in agreement with their induction by PMB, outer membrane localization and structural predictions.

## Discussion

While *arn* operon has been widely studied and is well known for its contribution to polymyxin resistance in *P. aeruginosa*; clinical isolates lacking the *arn* operon have been poorly reported. In this study, we have shown that a subgroup of isolates from respiratory and urine samples, lacks the entire *arn* operon. Interestingly, although *arn* operon is required to acquire stable resistance to polymyxins, adaptive resistance to polymyxins persisted in those strains suggesting that other genetic determinants were involved.

Exposure to sub-inhibitory concentrations of PMB in the *P. aeruginosa* IHMA87 strain strongly induced the expression of *mipBA* operon through a TCS, the inner membrane HK, ParS and the RR, ParR. At least three TCS (PmrAB, ParRS and CprRS) participate in the response to polymyxins in *P. aeruginosa* with partially overlapping regulons [9,12]. But ParR was the only response regulator binding to *mipBA* promoter in all three strains tested (PAO1, PA14 and IHMA87; [32]).

MipB and MipA proteins are associated with the membrane and form a complex *in vitro*. MipA localization in *Caulobacter crescentu*s was dependent on MreC, a proposed scaffold protein in bacterial elongasome [56,57]. Moreover, MipA was proposed to be tethered to both outer and inner membrane through interaction with MltA and PBP1b [58]. In *P. aeruginosa*, MipA appears to be embedded in the outer membrane, in line with its fold in β-barrel containing 12 β-strands that superimposes on the NanC and OmpG porins [47,48]. Structural predictions showed that MipA harbors a negatively charged inner channel in its center where several conformations of PMB and PME molecules can be docked, reminiscent to the crystal structure of the LSD1-CoREST bound to PMB (PDB entries: 5L3F and 5L3G; [55]). Further structural studies are essential to reveal the binding mode of these ligands to MipA. The MipA fold is observed in other proteins of Gram-negative bacteria suggesting that other outer membrane porins could bind positively charged molecules such as PMB or PME if the lateral gate and barrel lumen forming a negatively charged channel are conserved.

In light of structural characteristics of Mip proteins, and the finding that the mipBA deletion phenocopies the *parRS* mutant phenotype in presence of PMB, we propose that PMB binding to MipA modifies MipA/MipB structure and initiate the downstream signaling through the kinase activity of ParS, leading to the activation of ParR/S TCS, in turn allowing the induction of the MeXY-OprA pump and the adaptive resistance to PMB (model presented in Fig. 8). ParS is a classical HK composed of two transmembrane α-helixes, a 102 amino-acid sensing periplasmic domain and a cytoplasmic kinase domain. ParS belongs to the family of HKs sensing cAMPs, however how the sensing occurs at the molecular level is unknown. There is scarce structural information concerning the recognition of and signaling initiation by HK, probably due to their size, membrane localization and dynamics of the phosphate transfer (for review [59]). Recent high resolution structure of a nitrate/nitrite sensing histidine kinase, NarQ, revealed how the mechanistic signal can be amplified and propagated through the protein leading to kinase activity [60]. For example, it is established that the HK PhoQ of *Salmonella* is activated by cAMPs through displacement of magnesium ions (Mg^2+^;[61]). Therefore, small conformational changes in the perception domains of the sensing protein may lead to adapted transcriptional response.

**Figure 8.**
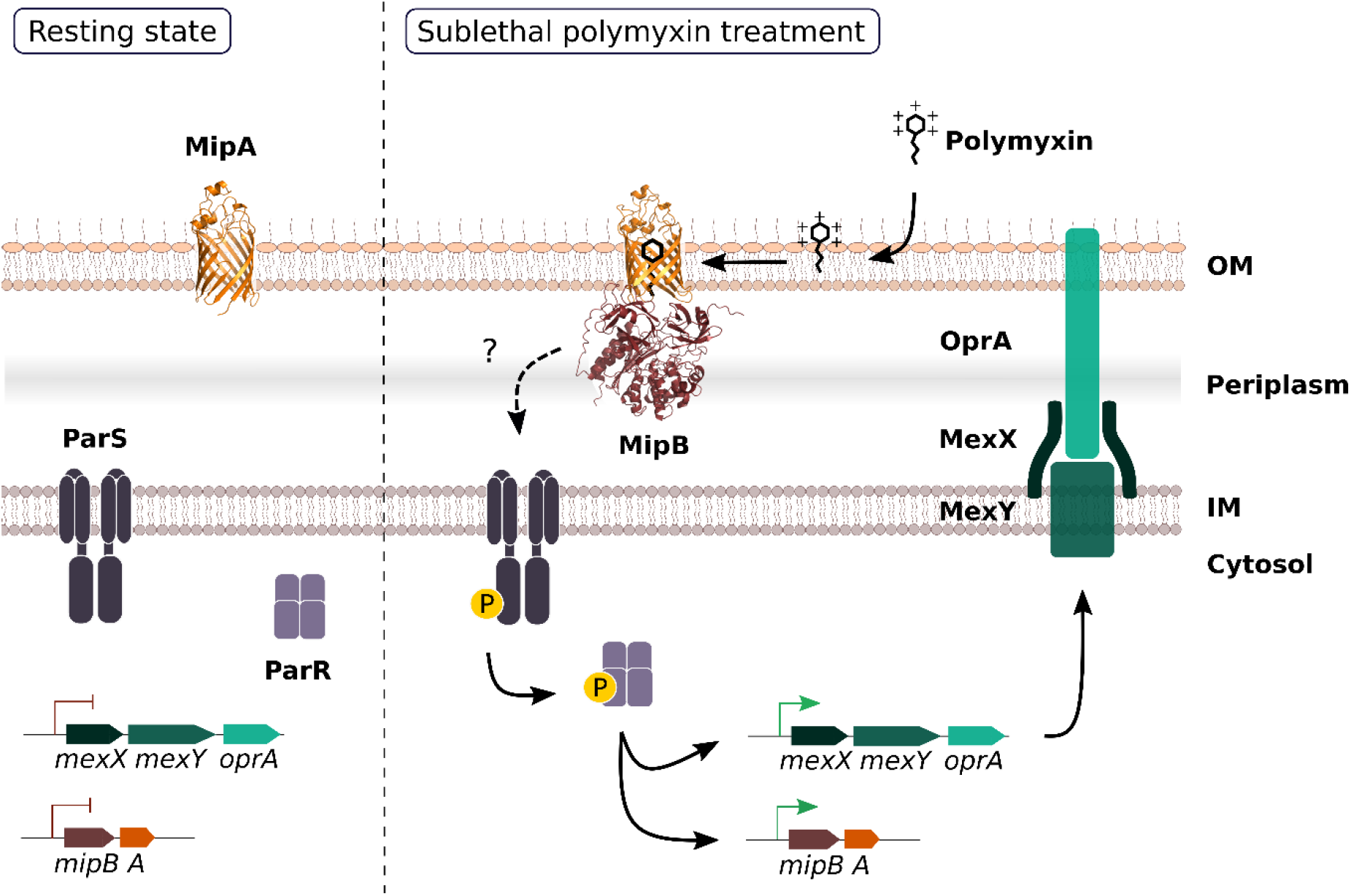
Schematic representation of a working model proposing MipA/MipB as co-sensors of PMB. In the resting state, low amounts of MipA are present in the outer membrane, and ParR/ParS TCS is inactive. PMB binding to bacterial membranes and to MipA provokes conformational changes in MipB-MipA complex. MipA and/or MipB induce ParS auto-phosphorylation which leads to activation of the cognate response regulator ParR. ParR binds to promoter regions of *mexXY-oprA* and *mipBA* operons resulting in MipA, MipB and MeXY-OprA overproduction and adaptive resistance to polymyxins.

Co-sensing or signal transfer between macromolecules occur in several signal transduction TCSs in Gram-positive bacteria (for review [62]). The most studied system is the *Bacillus subtilis* Bce module providing antimicrobial peptide resistance. This stress envelope response module is composed of an ABC transporter, BceA/BceB, which forms a complex with the inner membrane sensor kinase, BceS, of the TCS BceS/BceR [63,64]. Architecture of a complete Bce module by cryo-EM revealed extensive structural flexibility of the kinase BceS upon the bacitracin-dependent ATP binding to the ABC transporter, BceAB [65].

Moreover, MipB/MipA - ParR/ParS module shares striking functional and structural analogy with *E. coli* surface sensing module composed of an outer membrane lipoprotein NlpE that interacts with OmpA [66] acting as a membrane sensor of the envelope stress response TCS CpxR/CpxA [67]. The flexible N-terminal domain of NlpE interacts directly with the periplasmic domain of the CpxA kinase [68,69] to initiate the downstream phosphorelay. Indeed, using PDBeFold [45], CTD2 of MipB was superimposed onto the β-barrel M21-M99 of the sensor liporotein NlpE with a rmsd value of 2.22 Å. Investigating detailed molecular interactions between MipA/MipB and ParS will be a challenge of our future work.

Using the *arn* operon as a read-out, Fernandez et al., showed that ParS/ParR operon in *P. aeruginosa* responds to a variety of cAMPs, with the best inducer being indolicidin, followed by PMB [9]. They also showed that the inactivation of *mipB* by transposon insertion in PAO1 lead to reduced adaptive resistance to PMB [9]; however how the signaling occurs without the complete MipA is difficult to apprehend, although we cannot exclude that structural homologues of MipA exist in the PAO1 group of strains.

The study of *mipA* genetic environment across Gram-negative bacteria performed by WebFlaG [70] highlighted the presence of MipA-like outer membrane proteins in many distinct species; however, its genetic association with *mipB* is only present in species from the *Pseudomonas* genus (Genetic neighbor 3, Fig. S7). It is worth noting that a gene coding for another shorter serine hydrolase domain-containing protein was found next to *mipA* in the *Achromobacter* genus. Interestingly, the screen of *mipA* loci revealed that, in most cases and even in absence of *mipB*, a TCS was encoded just upstream or downstream of *mipA*, further suggesting a role of MipA in signaling and/or stress response (Genetic neighbor 1 and 2, Fig. S7).

Interestingly, in the strain devoid of the *arn* locus, the only membrane proteins induced by PMB were MexX, MexY and OprA, forming the resistance-nodulation (RND) cell division type efflux pump MexXY-OprA [71]. The outer protein OprA is lacking in PAO1 genetic background, and the periplasmic MexX protein, and cytoplasmic membrane protein MexY function in cooperation with the outer membrane protein OprM [72]. This active efflux pump has a wide substrate specificity (but not polymyxins) and contributes to intrinsic and acquired resistance to aminoglycosides in *P. aeruginosa* strains [73]. MexXY-overproduction is very frequent in clinical strains from health-associated infections and cystic fibrosis [74,75]. *mexXY* belongs to the ParR regulon and together with ParR/ParS-upregulated genes of the polyamine biosynthetic pathway (PA4773-PA4774-PA4775), contribute to PME/colistin tolerance and acquired resistance [76]. The molecular mechanism leading to PMB protection by MeXY-OprA is still unknown, and the IHMA87 strain represents the opportunity to investigate this aspect in the *arn*-negative genetic background.

Altogether, we provide new insights into *P. aeruginosa* global response to polymyxins and the heterogeneity among *P. aeruginosa* strains. We propose that MipB-MipA constitute a new sensor of polymyxins, which may signal outer membrane perturbation to HK ParS, of the ParR/ParS two-component signaling system, to orchestrate the bacterial response to PMB and probably other antimicrobials targeting outer membranes.

## Material and methods

### Bacterial strains and genetic manipulations

*E. coli* and *P. aeruginosa* were grown in Lysogeny Broth (LB) at 37°C under agitation. *P. aeruginosa* was selected on irgasan (25 µg/mL)-containing LB plates. Antibiotic were used at the following concentration: 75 µg/mL gentamicin, 75 µg/mL tetracycline and 100 µg/mL carbenicillin for *P. aeruginosa* and 50 µg/mL gentamicin, 10 µg/mL chloramphenicol, 100 µg/mL ampicillin for *E. coli.* Unless specified otherwise, *s*ub-lethal concentration of PMB used for *P. aeruginosa* strains was 0.25µg/mL.

All strains and plasmid are listed in Table S2.

Plasmid for protein expression pET15b-VP-*mipB-Strep/mipA-6His* and pET15b-VP-*mipA-6His*, were constructed first by amplifying the operon *mipBA* and the gene *mipA,* respectively by PCR and cloned by Sequence- and Ligation-free cloning (SLIC - [77]) reaction into the pET15b-VP vector (S. Lory lab). Of note, the pET15b-VP is a vector designed for expression in both *E. coli* and *Pseudomonas* sp. thanks to two specific *ori* sites. During the PCR reaction we add the sequence for six histidine residues at the end of the *mipA* gene. Then the pET15b-VP-*mipB/mipA-*6His was mutated in order to add a Strep-tag at the C-terminus of MipB, the Strep-tag sequence was optimized using *Pseudomonas* codon usage (TGGAGCCACCCGCAGTTCGAAAAG). The resulting vector was verified by sequencing. All primers were listed in Table S3.

For gene deletion, allelic exchange vectors were designed with upstream and downstream flanking regions of approximately 500 bp and cloned into a pUC57 (Amp) with *Hin*dIII and *Eco*RI (Genewiz). Fragments for *mipA*, or *mipBA* deletions were subcloned into pEXG2 (Gm). The allelic exchange vectors were conjugated into *P. aeruginosa* by triparental mating, using pRK600 as a helper plasmid. Clones resulting from homologous recombination were selected on irgasan containing LB plates and streaked onto NaCl-free MB plates with 10% sucrose (w/v) to select for plasmid loss. Sucrose resistant mutants were verified for the gene deletion by PCR after verification of antibiotic sensitivity.

### Phylogenetic analysis

144 *P. aeruginosa* genomes were retrieved from the Refseq database on the NCBI platform. New annotations were generated when needed. Altogether, the database included 8 genomes from phylogenetic group 1, 7 from group 2, 68 from group 4, 38 from group 5 and 23 from group 3. This database was remodeled with the addition of clinical strains sequenced by the French National Reference Center for Antibiotic Resistance, Besançon. Distances between genomes were established using Mash v2.3 [78] and a phylogeny was generated with mashtree v1.2.0 [79]. Mashtree uses any common sequence file and calls the neighbor-joining (NJ) algorithm which is implemented in the software QuickTree [80].

Search for genes of interest was performed using the Sequence Extractor plugin of BioNumerics 7.6.1 (Biomérieux) with at least 80 % homology and 90 % coverage to reference genes.

### Susceptibility testing

The Minimum inhibitory concentrations (MIC) of selected antibiotics were determined by broth microdilution method in Mueller Hinton broth (MHB, Becton Dickinson) with adjusted concentrations of Mg^2+^ (from 10 to 12 µg/mL), and Ca^2+^ (from 20 to 25 µg/mL) in agreement with CLSI recommendations [81].

Isolation of one-step mutants with stable colistin/PME resistance was performed by plating 100 µL aliquots of log phase *P. aeruginosa* PAO1, PA14, PA7 and IHMA87 cultures (A_600 nm_ equal to 1) on MHA supplemented with 4 to 64 µg/mL of colistin.

Drug bactericidal activity. Overnight cultures of PAO1, PA14, PA7 and IHMA87 were diluted into fresh pre-warmed MHB to yield an absorbance of *A*_600nm_ = 0.5 ± 0.05. The bacteria were incubated with a constant shaking (250 rpm) at 35°C for 30 min prior to the addition of colistin at a final concentration ranging from 0.5 to 16 µg /mL. 50 µL of the culture were transferred in a sterile tube at selected time point and inoculated on MH agar plate using easySpiral Spiral plater system (VWR). The survivors were counted after an overnight culture. Results were expressed as means of at least three independent experiments.

Stepwise adaptation of *P. aeruginosa* to colistin was obtained by increasing the drug concentration gradually in MHB. After a first exposure to 1/2 x MIC for 12 h, the bacteria were centrifuged and resuspended twice in drug-free medium prior to treatment with 2 x MIC for an additional 12 h period of time. Sequential exposures were carried out up to 64 x MIC.

### ParR purification

From genomic DNA of PAO1 reference strain, a fragment of 705 bp corresponding to the complete coding sequence of the gene *parR* without the TGA stop codon, replaced by XhoI restriction site, was amplified with specific primers PR1f (5’-GGTGAATTCATGGACTGCCCTA-3’) and PR1r (5’-CTCCTCGAGGAGCTCCCAGCCCAG-3’). It was subsequently cloned into the pET28(a) vector using the restriction enzymes *Eco*RI and *Xho*I to generate the plasmid pET98 (pET28ΩparR). The pET98 recombinant plasmid was then transformed into *E. coli* BL21λDE3/pREP4 [82] and transformants were selected on ampicillin (100 µL/mL). The transformants were grown in one liter of LB medium with constant shaking at 110 rpm at 30°C until an OD_600_=0.8 before adding 1 mM of IPTG (for 4 hours) to induce the overproduction of the ParR protein. After centrifugation and sonication of the bacterial pellet in ice, crude protein extract was loaded on 1-mL His trap fast-flow column (GE, Healthcare) equilibrated with buffer A (50 mM NaH_2_PO_4_, pH 7.5, 300 mM NaCl, 30 mM imidazole), and the protein was eluted with an imidazole gradient (300 mM-500 mM) using the AKTA prime chromatography system (GE Healthcare). Fractions containing pure ParR protein (26.5 kDa) after visualization on a Coomasie gel (14%) were pooled and dialysed against buffer B (50 mM NaH_2_PO_4_, pH 7.5, 300 mM NaCl, 50% glycerol), prior to determining protein concentration using the Bio-Rad Protein Assay.

### EMSA

From the genomic DNA of strain PAO1, the intergenic region (103 bp) of the *parR* and *mipB* genes containing the promoting region of genes *mipBA* (P*_mipBA_*) was labeled by PCR amplification using a combination of unlabeled primer with a primer end-labeled (625 nM) PM (5’-GACCCCGTTGACAGCG-3’) and PMrev (5’-TGGAACACCTGGCGGAAA-3’) with T4 polynucleotide kinase (0.075 U/µL) (New England Biolabs) and [g^32^P]-ATP (3000 Ci/mmol) (Perkin Elmer). The amplication was carried out in a 50 µL volume and the products were purified as previously described [83]. In order to phosphorylate the ParR protein, 150 µM of ParR protein was incubated in 20 µL of buffer C (50 mM Tris-HCL, pH 7.8, 20 mM MgCl_2_, 0.1 mM dithiothreitol) containing 178 pmol of acetyl [^32^P] phosphate (Hartmann Analytical) at 30°C for 1.30 h (ParR-P). The reactional mixture was loaded on a Sephadex G-50 spin column equilibrated with buffer C to remove excess of acetyl [^32^P] phosphate. The purified P*_mipBA_* labeled probes were incubated with various concentrations of purified ParR unphosphoryled (ParR) and phosphorylated (ParR-P) at 30°C for 20 min in 20 µL of buffer C. Then, the mixture was loaded with the DNA dye solution (40 % glycerol, 0.025 % bromophenol blue and 0.025 xylene cyanol) on a 7.5 % polyacrylamide gel. The gels were dried and analyzed by autoradiography.

### β-galactosidase assays

Bacteria were grown to mid-exponential phase and PMB (0.25 µg/mL unless stated otherwise) and further grown for 90 min. β-galactosidase activity was assessed according to [84] as described in [85]. Briefly, 500 µL of bacteria were permeabilized by the addition of 20 µL of 0.1% SDS and 20 µL of chloroform and vortexing step of 1 min. 100 µL of permeabilized bacterial suspension was added to 900 µL of Z buffer (0.06 M Na_2_HPO_4_, 0.04 M NaH_2_PO_4_, 0.01 M KCl, 1 mM MgSO_4_, pH 7) supplemented with β-mercaptoethanol (0.27 % (v/v)) and incubated at 28°C. Enzymatic reaction was started by the addition of 200 µL of *ortho*-Nitrophenyl-β-galactoside (ONPG, 4 mg/mL) and stopped with 500 µL of 1 M Na_2_CO_3_ solution. Absorbance at 420 nm was read using a spectrophotometer after sedimentation of cell debris. Promoter activities were expressed in Miller units ((A_420nm_×1000)(*t*_min_ x Vol_mL_x OD_600nm_)). Experiments were performed in three biological replicates.

### Protein predictions and genetic neighbor analysis

Signal peptides were predicted using SignalP 5.0 [86].

The 3D structure predictions of MipB and MipA/MipB complex were performed using AlphaFold [39] and AlphaFold-Multimer [53]. Pymol was used to calculate electrostatic surface potential and generate figures. WebFlaG was used to study genetic neighbors of *mipA* [70]. MipA FASTA sequence from MipA-IHMA87 was imputed and a list of 50 homolog was searched by BlastP in the Atkinson lab reduced database.

### Bacterial fractionation

Bacterial fractionation was performed according to the protocol described in [87]. Briefly, bacteria were grown to exponential phase (OD_600nm_ of 1). A total amount of 10^10^ bacteria were pelleted at 4°C and resuspended in 1 mL of buffer A (20 mM Tris-HCl pH 8, 200 mM MgCl_2_) supplemented with protease inhibitor cocktail (PIC, Roche) and lysozyme to a final concentration of 0.5 mg/mL. The sample was incubated for 30 min at 4°C on a rotating wheel. Periplasmic fraction was recovered after centrifugation at 11,000 xg for 15 min at 4°C. The spheroplast-containing pellet was washed with 1 mL of buffer B (20 mM Tris-HCl pH 8, 20 % sucrose) supplemented with PIC and resuspended in 1 mL of buffer B before a sonication step (5 min, 40 % intensity, 10 s ON/10 s OFF) on ice. Bacteria debris were removed by centrifuging for 15 min at 8,000 xg at 4°C and supernatant, composed of bacterial cytosol and membranes was further centrifuged for 10 min at 6000 rpm at 4°C. Cytosolic and membranes components were separated by ultracentrifugation at 200,000 xg (Rotor TLA120 Beckman) for 45 min at 4°C. The supernatant containing the cytosol was recovered, the pellet was washed twice with 1 mL of buffer C (20 mM Tris-HCl pH 8.0, 20 mM MgCl_2_) supplemented with PIC. Bacterial membranes were resuspended in 50 0µL of buffer B using a potter. All fractions were resuspended or diluted in 4X SDS-PAGE loading buffer and boiled for 10 min prior to SDS-PAGE or Western blot analyzes.

### Inner and outer membrane separation

Inner and outer membranes were separated using a discontinuous sucrose gradient as described in [51]. In brief, bacteria were grown to exponential phase and 2.5 x 10^11^ bacteria were pelleted for 15 min at 4°C and resuspended in 25 mL of buffer A (10 mM Tris-HCl pH 7.4, RNase 10 µg/mL, DNase 10 µg/mL and 20 % sucrose) supplemented with PIC. Bacteria were lyzed by sonication at 50 % intensity, 7 min, 30 s ON/ 30 s OFF) and remaining cell debris were removed by a centrifugation step. Supernatants were ultracentrifuged at 200,000 xg for 45 min at 4°C (TI45 Beckman rotor). Total membrane pellet was resuspended in 500 µL of buffer B (10 mM Tris-HCl pH 7.4, 5 mM EDTA, 20 % sucrose and PIC) and loaded onto an eight-1.5mL-layers sucrose gradient (from bottom to top) with sucrose concentrations of 55 %, 50 %, 45 %, 40 %, 35 % and 30 %. Another ultracentrifuge step of 72 h at 90,000 xg (Beckman SW41 swinging rotor) at 4°C without brake. Finally, 500 µL fractions were collected and further characterized by SDS-PAGE and Western blot analyzes.

### Western blot analyses

Sample protein content present on the SDS-PAGE was transferred onto a polyvinylidene difluoride (PVDF) membrane (GE Healthcare) for 90 min (25 V, 125 mA) in Laemmli buffer with 20 % ethanol. Membranes were blocked for 1 h at room temperature in 5 % (w/v) dry milk in PBS-Tween 0.1 % and labeled with primary antibodies for 1 h. Primary antibodies were used at the following concentrations: anti-MipA (1/ 2,000, rabbit, Biotem), anti-DsbA (1/10,000, obtained from R. Voulhoux, CNRS, Marseille, France), anti-Ef-TU (1/10,000, mouse, Hycult biotech), anti-XcpY (1/2,000, rabbit, [88]), anti-Streptag (1/6,000, mouse) and anti-His_6_ (1/6,000, mouse). The secondary horseradish peroxidase (HRP)-conjugated antibodies directed against rabbit or mouse were used at a 1/20,000 dilution (Sigma). Detection of luminescent signal was performed with a Luminata Western HRP substrate kit (Millipore).

Polyclonal rabbit antibodies were raised against urea treated purified MipA-6His following the manufacturer recommendations (Biotem) and used for immunodetection.

### MipA/MipB complex purification

*E. coli* BL21(DE3)RIL strain containing the pET15b-VP-*mipB*-Strep/*mipA*-6His plasmid were grown in LB (1 L) with appropriate antibiotic concentrations at 37°C with shaking. When bacteria reached an OD_600nm_ of 0.5-0.7, protein expression was induced with 1 mM IPTG and further grown for 2.5 h at 37°C with shaking. Bacterial culture was then centrifuged at 6,000 xg, for 20 min at 4°C and pellet was resuspended in 40 mL of lysis buffer (100 mM Tris-HCl pH 8.0, 150 mM NaCl, 10 % glycerol (v/v), 1 mM EDTA and 2 % N-lauroylsarcosine (w/v)) supplemented with Protease Inhibitors Cocktail (cOmplete ULTRA Tablets Roche). Bacterial cells were lysed with a M110-P microfluidizer (Microfluidics) with three passages at 15,000 psi and centrifuged at 39,000 xg for 45 min at 4°C to remove bacterial debris. The soluble fraction was loaded onto a StrepTap HP 1 mL affinity column (GE Healthcare) at 4°C previously equilibrated with lysis buffer. The column was washed with 5 CV of lysis buffer before elution with the same buffer containing 2.5 mM of desthiobiotin (Sigma D1411). Eluted fractions were analyzed by 12 % SDS-PAGE stain by Coomassie Blue and immunoblotting with anti-His, and anti-MipA for MipA presence and anti-Strep for MipB presence.

### MipA purification

Freshly transformed *E. coli* BL21(DE3)C41 colonies harboring pET15b-VP-*mipA*-His_6_ were grown at 30°C overnight in LB with 100 µg/ml Ampicillin (Amp) with shaking. The next day culture was diluted in 1 L LB-Amp and grown at 37°C until an OD_600nm_ of 1.0, then protein expression was induced with 0.5 mM IPTG and further grown for 2.5 h at 37°C with shaking. Bacterial culture was then centrifuged at 6,000 xg for 30 min at 4°C and pellet was resuspended in 200 mL of lysis buffer (25 mM Tris HCl pH 8.0, 150 mM NaCl, 10 % glycerol (v/v), 2 % N-lauroylsarcosine (w/v)) supplemented with Protease Inhibitors Cocktail (cOmplete ULTRA Tablets, Roche), DNaseI (1 µg/ml) and RNaseI (10 µg/ml). Bacterial cells were lysed with a M110-P microfluidizer (Microfluidics) with ten passages at 15,000 psi and centrifuged at 39,000 xg for 1 h at 4°C to remove bacterial debris. The soluble fraction was loaded at low speed (0.2 ml/min) at 4°C onto a HisTrap-HP 1 mL affinity column (GE Healthcare) equilibrated with buffer A (20 mM NaPi pH 7.7, 500 mM NaCL, 20 mM imidazole and 1 % N-lauroylsarcosine). The column was washed with 40 volumes of buffer A before elution in with the same buffer A containing 175 mM imidazole. 10 mM DTT and 5 mM EDTA were added to the fractions containing MipA-His_6_ to avoid disulfide bridges formation and inhibit the metalloproteases. The protein samples were then injected onto a Superdex200 Increase 10/300 GL (GE Healthcare) equilibrated with 25 mM HEPES, pH 7.5, 150 mM NaCl, 1 mM EDTA and 0.096 % LAPAO (w/v). MipA-His_6_, eluted in the peak corresponding to elution volume (Ve) of 12 ml, was recovered and EDTA was removed by 4 washing steps to allow binding to His-Trap column. Protein was finally concentrated on affinity column equilibrated with 25 mM Tris-HCl pH 8.0, 150 mM NaCl, 0.1% LAPAO and eluted in one step of the same buffer containing 150 mM imidazole. Protein purity was analyzed by 15 % SDS-PAGE, and protein concentration was determined by OD_280_ nm.

### Protein stability measurement

The protein stability and conformational changes were determined using differential scanning fluorimetry (DSF) that allow to use dye-free samples containing detergent. Capillaries were filled by samples and loaded on the Prometheus NT48 using nano-DSF technology (Nanotemper). Temperature was increased 1°C/min from 20°C to 95°C and fluorescence at 330 nm (F330) and 350 nm (F350) was recorded. The ratio (F350/F330) was calculated and plotted. The inflection point corresponding to a maximum in the first derivative (slope) of F350/F330 give the value of the melting temperature (Tm).

### MS-based quantitative proteomic analysis of bacterial membranes

Bacterial cultures (30 mL LB) were inoculated at an OD_600nm_ of 0.1 and grown to OD_600nm_ ∼ 0.6. PMB was added when needed (0.25 µg/mL) for 90min. Bacteria were harvested and resuspended in 1 mL of buffer B (same buffers as in bacterial fractionation) supplemented with PIC. Bacteria were lyzed by sonication during 5 min (40 % intensity, 10s ON/10 s OFF) and bacterial debris were eliminated by a centrifugation step of 15 min at 6,000xg at 4°C. Supernatant was ultracentrifuged at 200,000 xg for 45 min at 4°C and soluble fraction was removed. Membranes pellet were washed, resuspended in 100 µL of buffer C and loading buffer was added and samples were boiled for 10 min. Presence of MipB and MipA in each sample was assessed by western blot analysis.

Three replicates of membrane fraction were prepared for each analysed strain. The proteins were solubilized in Laemmli buffer and stacked in the top of a 4-12% NuPAGE gel (Invitrogen). After staining with R-250 Coomassie Blue (Biorad), proteins were digested in-gel using modified trypsin (sequencing purity, Promega), as previously described [51]. The resulting peptides were analyzed by online nanoliquid chromatography coupled to MS/MS (Ultimate 3000 RSLCnano and Q-Exactive HF, Thermofisher Scientific) using a 180 min gradient. For this purpose, the peptides were sampled on a precolumn (300 μm x 5 mm PepMap C18, Thermo Scientific) and separated in a 75 μm x 250 mm C18 column (Reprosil-Pur 120 C18-AQ, 1.9 μm, Dr. Maisch). The MS and MS/MS data were acquired using Xcalibur (Thermo Fisher Scientific).

Peptides and proteins were identified by Mascot (Matrix Science) through concomitant searches against the NCBI database (*P. aeruginosa* strain: IHMA879472 taxonomy, March 2021 download) and a homemade database containing the sequences of classical contaminant proteins found in proteomic analyses (human keratins, trypsin, etc.). Trypsin/P was chosen as the enzyme and three missed cleavages were allowed. Precursor and fragment mass error tolerances were set at respectively at 10 and 20 ppm. Peptide modifications allowed during the search were: Carbamidomethyl (C, fixed), Acetyl (Protein N-term, variable) and Oxidation (M, variable). The Proline software version 2.2.0 [89] was used for the compilation, grouping, and filtering of the results (conservation of rank 1 peptides, peptide length ≥ 6 amino acids, false discovery rate of peptide-spectrum-match identifications < 1% [90], and minimum of one specific peptide per protein group). Proline was then used to perform a MS1 label-free quantification of the identified protein groups based on razor and specific peptides.

Statistical analysis was then performed using the ProStaR software version 1.30.5 [91]. Proteins identified in the contaminant database, proteins identified by MS/MS in less than two replicates of one condition, and proteins detected in less than three replicates of one condition were discarded. After Log_2_ transformation, abundance values were normalized by vsn (variance stabilizing normalization) method before missing value imputation (slsa algorithm for partially observed values in the condition and DetQuantile algorithm for totally absent values in the condition). Statistical testing was then conducted using Limma, whereby differentially expressed proteins were sorted out using a Log_2_ (fold change) cut-off of 1 and a p-value cut-off of 0.01, leading to FDRs inferior to 3% according to the Benjamini-Hochberg estimator. Proteins found differentially abundant but identified by MS/MS in less than two replicates and detected in less than three replicates in the condition in which they were found to be more abundant were manually invalidated (*p*-value = 1).

### RT-qPCR

After total RNA extraction from 2 ml of bacterial cultures grown to OD_600_=1 using TRIzol™ Plus RNA Purification Kit and Turbo DNAse treatment, cDNA synthesis was carried out with the SuperScript IV Reverse Transcriptase (Invitrogen). The absence of genomic DNA was verified for each sample. Experiments were performed with 4 biological replicates for each strain and condition, and in technical duplicate. Quantification is based on real-time SYBR green amplification molecules with specific target primers using Luna® Universal qPCR Master Mix (Biolabs). Relative mRNA expression is calculated using the CFX Manager software (Bio-Rad), with *rpoD* reference gene quantification. Statistical analyses were performed with Graphpad prism 9. The sequences of primers are listed in supplementary table S3.

### Statistical analysis

Data were statistically analyzed using Graphpad prism 9. For multiple comparisons, a one-way ANOVA or Kruskall-Wallis test was performed depending on the data normality (Shapiro test), followed by a Tukey or a Dunn’s test for normally and non-normally distributed dataset, respectively. To compare two groups, a Student t-test or a Mann-Whitney test was applied depending on the normality of the data.

### Data availability

MS data have been deposited to the ProteomeXchange Consortium via the PRIDE partner repository [92] with the dataset identifier PXD043253. Tables are available upon request.

## Acknowledgements

This work was supported by grants from Agence Nationale de la Recherche (ANR-18-CE11-0019) and from GRAL, funded through the University Grenoble, Alpes graduate school (Ecoles Universitaires de Recherche) CBH-EUR-GS (ANR-17-EURE-0003). *P. aeruginosa* IHMA87 was obtained from the International Health Management Association, USA. MJM received a Ph.D fellowship from the French Ministry of Education and Research. The proteomic experiments were partially supported by Agence Nationale de la Recherche under projects ProFI (Proteomics French Infrastructure, ANR-10-INBS-08) and GRAL, a program from the Chemistry Biology Health (CBH) Graduate School of University Grenoble Alpes (ANR-17-EURE-0003). We also thank undergraduate students Claire Lapouge and Florine Borrel for their technical help during experiments.

