## Supplemental information for "*Pseudomonas aeruginosa* MipA-MipB envelope proteins act as new sensors of polymyxin"

### Supplementary information

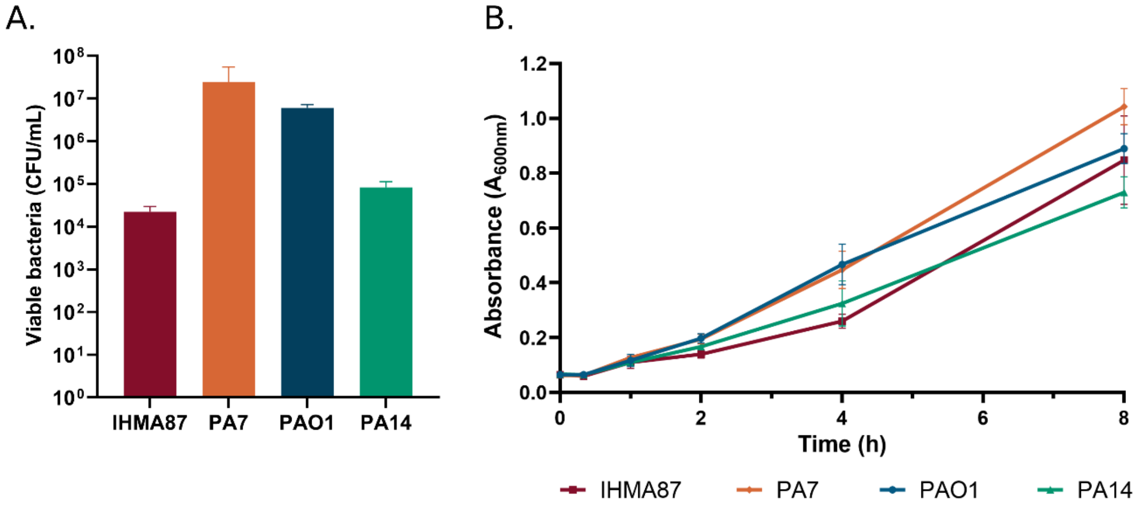

**Figure S1. *Arn* negative strains are capable of adaptation to polymyxins. A.** Bacterial tolerance to PME. Number of viable bacteria at 64µg/mL of PME. n=3. **B.** Bacterial growth curve. Strains were grown and absorbance at 600nm was measured at different timepoints. n=3.

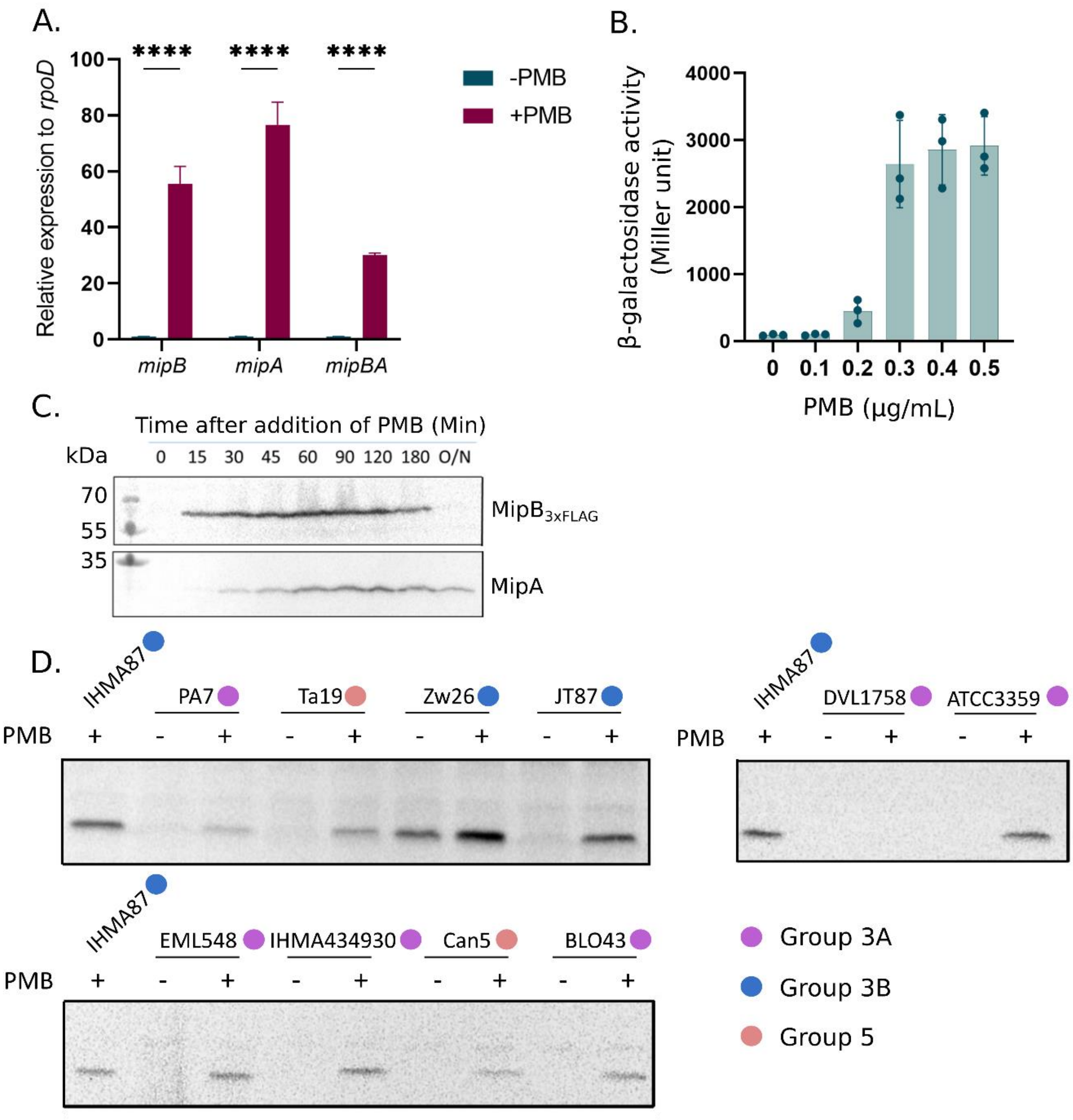

**Figure S2. *mipBA* induction upon sub-lethal PMB treatment is rapid, dose-dependent and conserved among *P. aeruginosa* strains. A.** *mipBA* operon is induced in reponse to sublethal concentration of PMB measured by RT-qPCR. n=3. **B.** Dose-response of $P_{mipBA}$ promoter activity to PMB performed by β-galactosidase assay. n=3. **C.** Kinetics of MipB and MipA synthesis upon PMB sublethal treatment. **D.** The induction of MipA in response to PMB is conserved across strains from group 3A, groups 3B and 5 in *P. aeruginosa* species.

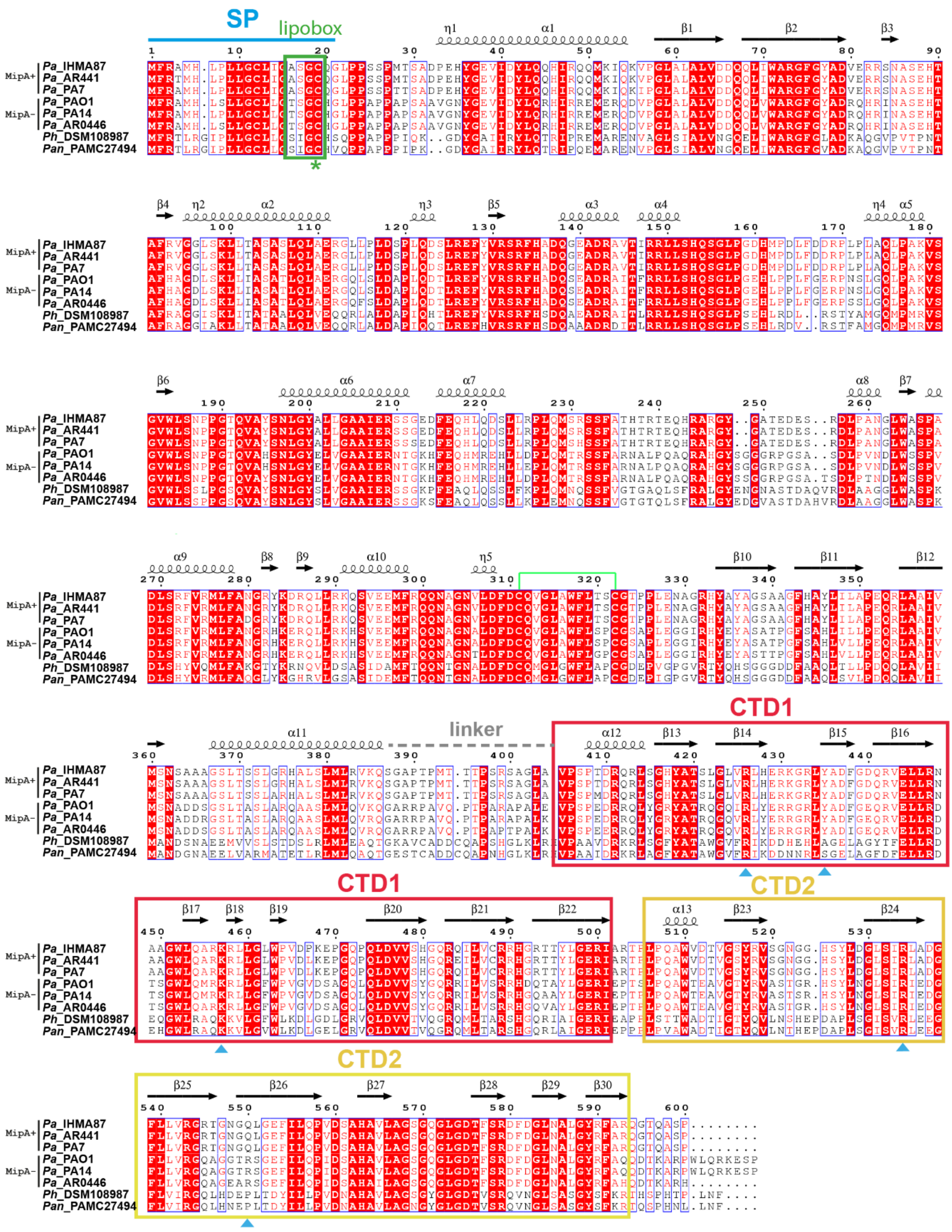

**Figure S3. Sequence alignment of MipB proteins.** Proteins homologous to *P. aeruginosa* MipB from *P. aeruginosa* IHMA87 were aligned using CLUSTAL. The secondary structure of MipB-IHMA87 retrieved from the model is shown above the sequence alignment. The putative disulfide bridge between Cys311 and Cys321 is depicted with the green line. Note the divergent sequence between MipA+ and MipA-negative strains within the *P. aeruginosa* group. Strains used for alignment were IHMA87, AR441 and PA7 which contain MipA and PAO1, PA7 and AR0446 which only have the truncated version of MipA*. The MipB sequences of *P. heamolytica* strain DSM108987 and *P. antartica* PAMC 27494 were included in the alignment. Full and weakly conserved residues are shaded in red and blue boxes, respectively. MipB has a predicted lipobox (green box) with one Cys (asterisk). The predicted signal peptide

(SP), the large beta-lactamase like domain followed by the linker loop (dashed gray line) and the two C-terminal domains (CTD1 and CTD2) are shown. Note the high conservation of the CTDs inside the MipA+ and MipA- groups. The residues important for MipA interaction are highlighted by a cyan triangle. The alignment was visualized by ESPript 3 [1].

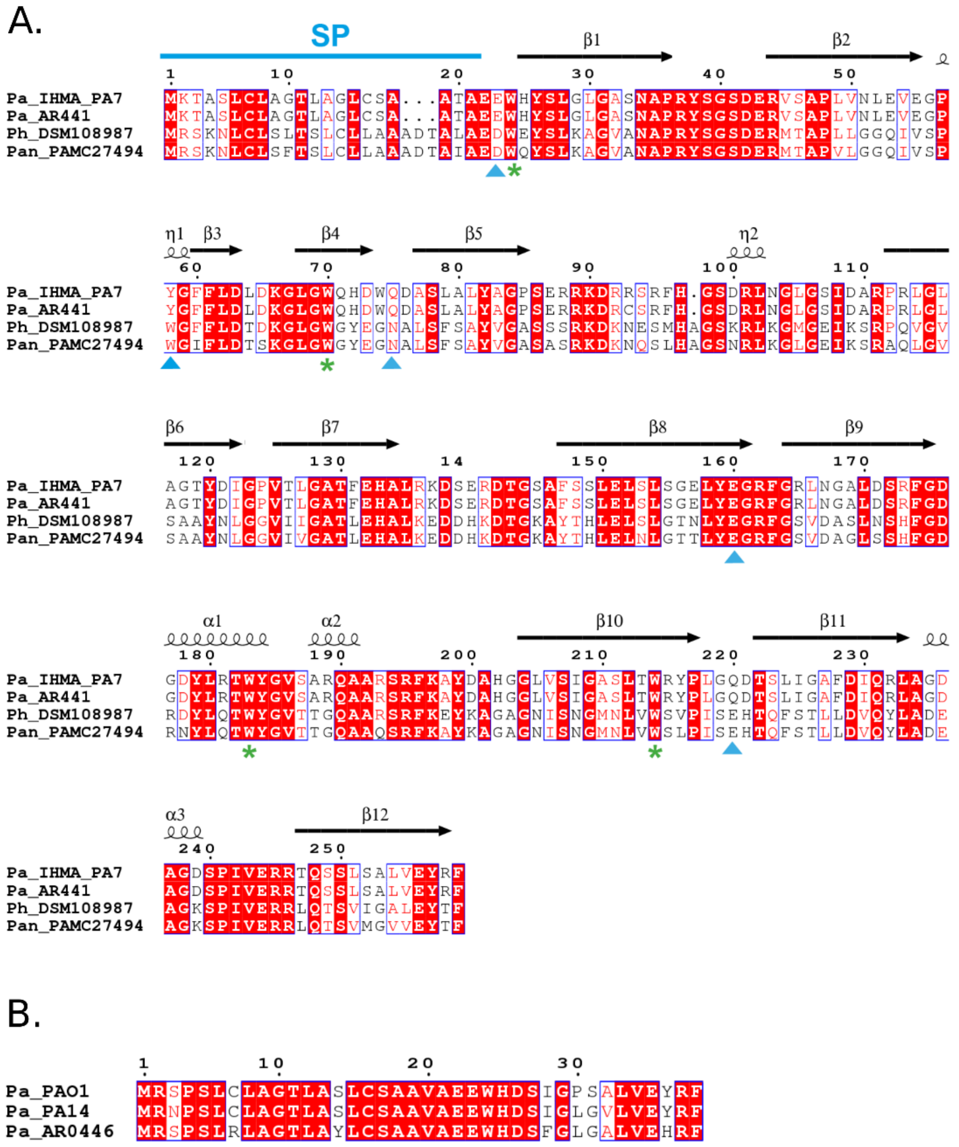

**Figure S4. Sequence alignment of MipA. A.** Proteins homologous to *P. aeruginosa* MipA of IHMA87 were aligned using CLUSTAL. The secondary structure of MipA-IHMA87 retrieved from the model is shown above the sequence alignment. Strains used for the alignment were *P. aeruginosa* IHMA87, PA7, AR441 and CR1, and *P. heamolytica* DSM108987 and *P. antartica* PAMC 27494. Full and weakly conserved residues are shaded in red box and blue box, respectively. The four-conserved Trp residues important for the OM insertion are highlighted by an asterisk, the residues important for interaction with MipB are depicted by a cyan arrow. Signal peptide (SP). The alignment was visualized by ESPript 3 [1]. **B.** Sequence alignment of short versions of the MipA protein from PAO1, PA14 and AR0446.

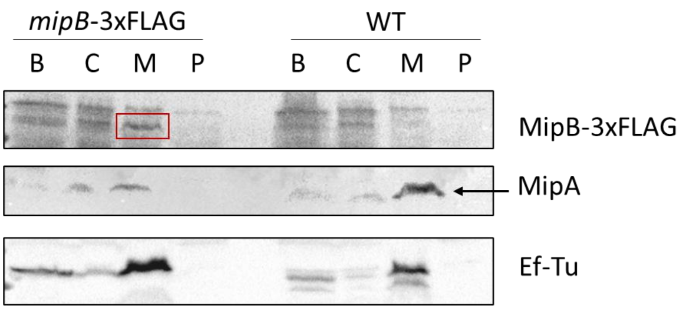

**Figure S5. MipB localizes in bacterial membranes.** Bacterial fractionation showing membrane association of MipA and MipB-3xFLAG. EF-Tu is a control for the cytosolic fraction. Lines B: whole bacteria, C: cytosol, M: total membranes, P: periplasm.

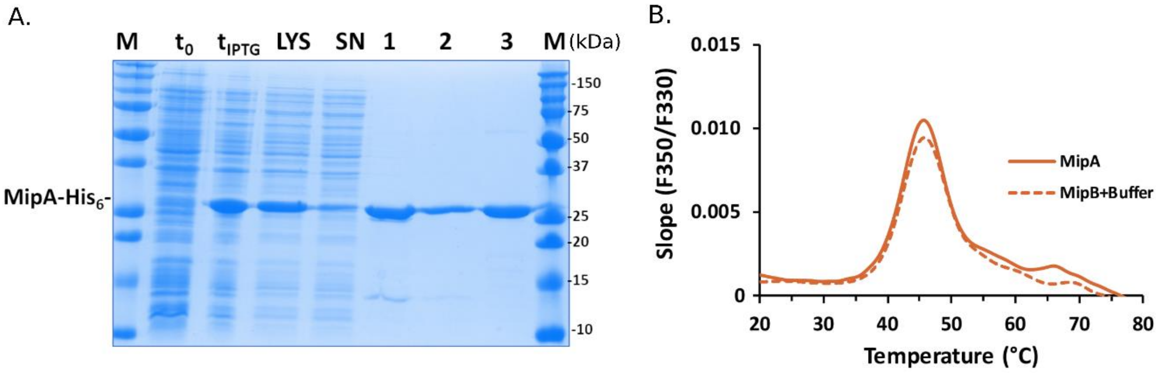

**Figure S6. Purified MipA was used for nanoDSF. A.** SDS-PAGE summarizing the purification steps of MipA-6His. Protein expression in *E.coli* was induced in exponential phase (t0) by IPTG addition ($t_{IPTG}$), bacteria were lysed (LYS) in presence of 2% N-lauroylsarcosine (w/v), the soluble proteins (SN) were loaded on an affinity column His-Trap and eluted with imidazole (**1**). MipA was further purified on Superedex200 (**2**) and concentrated on HisTrap (**3**) in presence of 0.1% LAPAO. **B.** The melting temperature of pure MipA (full line) was determined by differential scanning fluorimetry (DSF) by heating the samples from 20 to 95°C. As negative control the same amount of buffer without polymyxins was added to the sample containing MipA (dashed line).

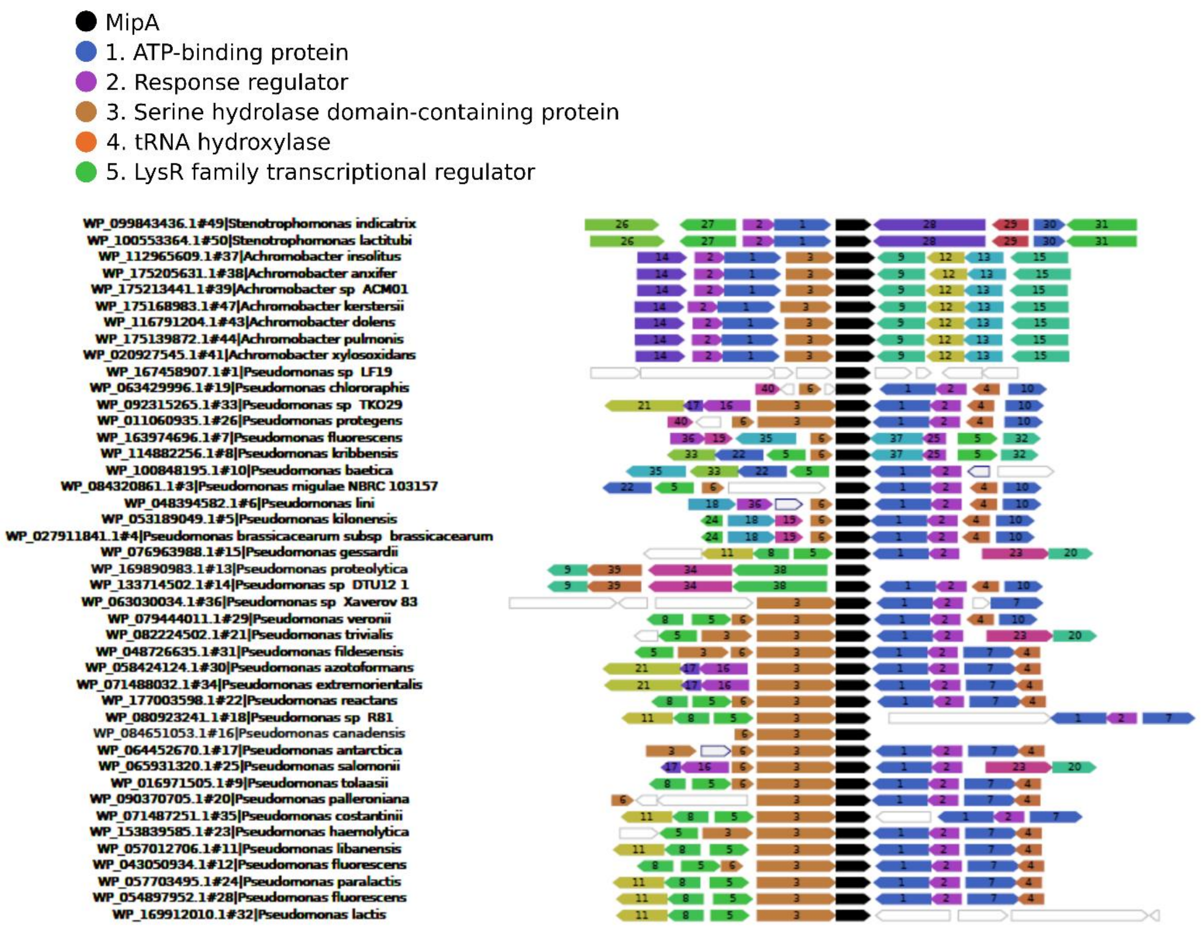

**Figure S7. *mipA* genetic neighborhood conservation.** 50 MipA protein homologs were retrieved by BLASP together with the corresponding neighboring genes within the Atkinson lab reduced database and most common genetic neighbors' protein predictions are indicated. Analysis and image generated with webFlaGs [2].

**Table S1. Data from proteomic analysis of bacterial membranes**

**Table S2. Bacterial strains and plasmids**

| Bacteria | Features / Source | Reference/origin |
|---|---|---|
| ***Pseudomonas aeruginosa*** | | |
| IHMA879472/ AZPAE15042 | Isolated from a urinary infection in Germany, group 3B | IHMA[1] collection [3,4] |
| IHMA87Δ*mipA* | IHMA87 *mipA* deletion mutant | This work |
| IHMA87Δ*mipBA* | IHMA87 *mipBA* deletion mutant | This work |
| IHMA87Δ*parRS* | IHMA87 *parRS* deletion mutant | This work |
| IHMA87::pminiCTX-*PmipBA*-lacZ | IHMA87 *with PmipBA-lacZ* transcriptional fusion ($Tc^R$) | This work |
| PA7 | Isolated from a wound in Argentina, group 3A | [5] |
| PAO1 | Isolated from a wound isolated, group 1 | [6], Lab collection |
| PA14 | Isolated from a burnt infection, group 2 | [7] |

| | | |
|---|---|---|
| EML548 | Isolate from Germany, group 3A | [8] |
| ATCC33359/ EML545 | Isolated from a water sample in Germany, group 3A | [8] |
| IHMA434930/ AZPAE14901 | Isolated from intra-abdominal tract infection in India, group 3A | IHMA[1] collection [8] |
| BL043 | Isolated from a bacteremia in the USA, group 3A | [8] |
| Zw26 | Isolated from a cystic fibrosis sputum in Germany, group 3B | [8] |
| JT87 | Isolated from a urinary infection in the USA, group 3B | [8] |
| Ta19 | Isolated from a urine sample in Australia, group 5 | [8] |
| Can5 | Isolated from a dog infection in the United Kingdom, group 5 | [8] |
| DVL1758 | Isolated from a shallow pond in Belgium, group 5 | [8] |
| ***Escherichia coli*** | | |
| DH5α | Laboratory strain | Lab collection |
| TOP10 | Cloning strain | Invitrogen |
| BL21(DE3)RIL | *E. coli* expressing extra copy of tRNA genes for codone rare: *arg*U (AGA, AGG), *ile*Y (AUA) and *leu*W (CUA) (Cm$^R$) | This work |
| BL21(DE3)C41 | *E.coli* for expression of MipA-His$_6$ | This work |
| BL21(DE3)RIL pET15bVP-*mipB*-*strep*/*mipA*-His$_6$ | *E. coli* for co-expression of MipA-His$_6$ and MipB-Strep | This work |
| **Plasmids** | | |
| pRK600 | Helper plasmid with conjugative properties (Cm$^R$) | [9] |
| pEXG2 | Allelic exchange vector (Gm$^R$), *sacB* | [10] |
| pEXG2-mut-*mipBA* | pEXG2 carrying DNA fragment for *mipBA* deletion in IHMA87 obtained by SLIC (Gm$^R$) | This work |
| pET15b-VP | Engeneered vector with two *ori* sites for expression of proteins in both *E.coli* and *Pseudomonas* | S. Lory Lab |
| pET15b-VP-MipB/MipA-His$_6$ | Vector with operon *mipAB* cloned into *Nco*I/*Bam*HI sites (Amp$^R$) | This work |
| pET15b-VP-MipB-Strep/MipA-His$_6$ | Vector used for co-purification of MipB-Strep and MipA-His$_6$ (Amp$^R$) | This work |
| pET15b-VP-MipA-His$_6$ | Vector use for purification of MipA | This work |

[1] International Health Management Association, USA

**Table S3. Primers**

| Primers | Sequence (5’-3’) | Purpose |
|---|---|---|
| MipBStrep-rv | Ccagggcacccaggcatcaccc TGG AGC CAC CCG CAG TTCGAAAAG tgaaccaatcgaaaggaatccctc | Used for mutagenesis (add of Strep tag sequence to Cter MipB) |
| MipBStrep-fw | gagggattcctttcgattggttcaCTTTTCGAAC TGCGGGTGGCTCCAgggtgatgcctgggt gccctgG | Used for mutagenesis (add of Strep tag sequence to Cter MipB) |
| NcoI-MipB | tttaagaaggagatataccatgttccgcgcaatgcat ctcc | Used for cloning *mipAB* operon by SLIC reaction in pET15b-VP |
| MipB-BamHI-6His | gctttgttagcagccggatcc**tca**gtgatgatgatga tgatggggtgatgcctgggtgccc | Used for cloning *mipAB* operon by SLIC reaction in pET15b-VP |
| MipA-BamHI-6His | gctttgttagcagccggatcctcagtgatgatgatgat gatggaagcggtattccaccagcgcg | Used for cloning *mipA* operon by SLIC reaction in pET15b-VP |
| NcoIMipA | tttaagaaggagatataccatgaaaaccgcctccct gtg | Used for cloning *mipA* operon by SLIC reaction in pET15b-VP |
| qPCR-mipA-F | tatccccttggccaggacac | Used for RT-qPCR |
| qPCR-mipA-R | cgagctttgcgtacgacgtt | Used for RT-qPCR |
| qPCR-mipB_F | gttcgccaacggccggtaca | Used for RT-qPCR |
| qPCR-mipB_R | aggacgttgccggcgttctg | Used for RT-qPCR |
| qPCR-mexA_F | cagaaccgcctgaagatcgt | Used for RT-qPCR |
| qPCR-mexA_R | gccgctttctccacgtagat | Used for RT-qPCR |
| qPCR-mexB_F | atgcacatccaatggaccgg | Used for RT-qPCR |
| qPCR-mexB_R | cagcccagcaggaataggg | Used for RT-qPCR |
| qPCR-oprA_F | ggcaacaacagttcaccgac | Used for RT-qPCR |
| qPCR-oprA_R | aggttgcggttgtgctcc | Used for RT-qPCR |
| qPCR-rpoD_F | ctgccggaggatatttcaga | Used for RT-qPCR |
| qPCR-rpoD_R | atacgttgatccccatgtcg | Used for RT-qPCR |
